# Haplotype-based inference of the distribution of fitness effects

**DOI:** 10.1101/770966

**Authors:** Diego Ortega-Del Vecchyo, Kirk E. Lohmueller, John Novembre

## Abstract

Recent genome sequencing studies with large sample sizes in humans have discovered a vast quantity of low-frequency variants, providing an important source of information to analyze how selection is acting on human genetic variation. In order to estimate the strength of natural selection acting on low-frequency variants, we have developed a likelihood-based method that uses the lengths of pairwise identity-by-state between haplotypes carrying low-frequency variants. We show that in some non-equilibrium populations (such as those that have had recent population expansions) it is possible to distinguish between positive or negative selection acting on a set of variants. With our new framework, one can infer a fixed selection intensity acting on a set of variants at a particular frequency, or a distribution of selection coefficients for standing variants and new mutations. We apply our method to the *UK10K* phased haplotype dataset of 3,781 individuals and find a similar proportion of neutral, moderately deleterious, and deleterious variants compared to previous estimates made using the site frequency spectrum. We discuss several interpretations for this result, including that selective constraints have remained constant over time.

## Introduction

The distribution of fitness effects for new mutations (*DFE*) is one of the most important determinants of molecular evolution. The *DFE* is a probability distribution that quantifies the proportion of new mutations having a certain selection coefficient *s*, where *s* can take positive or negative values depending on whether the allele is under positive or negative selection. The *DFE* determines current levels of genetic variation, since the frequencies of the alleles under selection depend on their selection coefficient (Sawyer & Hartl 1992; Hartl *et al*. 1994; Bustamante *et al*. 2001), and alleles under selection change the genetic variation at linked sites due to the effects of linked selection (Maynard Smith & Haigh 1974; Charlesworth *et al*. 1993). The *DFE* is also a key feature in the evolution of complex phenotypic traits (Lohmueller 2014a; Simons *et al*. 2014; Mancuso *et al*. 2015), since the association between the selection coefficients and the effect of mutations on a complex trait is an important determinant of the genetic architecture of a trait (Eyre-Walker 2010). Due to the impact of the *DFE* on levels of genetic and phenotypic variation, properly inferring the *DFE* is essential to many fundamental problems such as validating predictions of the nearly neutral theory (Kimura & Crow 1964; Crow 1972; Ohta 1992), understanding changes in the deleterious segregating variation observed in different populations (Gazave *et al*. 2013; Lohmueller 2014b; Henn *et al*. 2015; Brandvain & Wright 2016; Gravel 2016; Simons & Sella 2016; Koch & Novembre 2017), elucidating the factors that influence changes on the *DFE* between species (Martin & Lenormand 2006; Charlesworth & Eyre-Walker 2007; Serohijos & Shakhnovich 2014; Tenaillon 2014; Rice *et al*. 2015; Huber *et al*. 2017), and inferring the amount of adaptive evolution between species (Gossmann *et al*. 2012; Galtier 2016; Zhen *et al*. 2018).

Broadly, two lines of research have been developed to infer a *DFE*. One is based on experimental approaches and the other one is based on the analysis of population genetic variation at putatively neutral and deleterious sites. The main experimental approaches taken with viruses, bacteria and yeast are site-directed mutagenesis experiments in target regions (Bataillon & Bailey 2014) and mutation-accumulation experiments (Halligan & Keightley 2009). They are useful because they can obtain information about the *DFE* including advantageous and deleterious mutations; that said, advantageous mutations tend to be rare or not found in results from experimental approaches (Halligan & Keightley 2009; Lind *et al*. 2010; Jacquier *et al*. 2013; Bataillon & Bailey 2014) with some exceptions (Sanjuán *et al*. 2004; Dickinson 2008). The types of probability distributions that have provided a good fit to the *DFE* of deleterious mutations on site-directed mutagenesis experiments are a gamma distribution (Domingo-Calap *et al*. 2009; Lind *et al*. 2010; Jacquier *et al*. 2013), a unimodal distribution with a similar shape to a gamma distribution (Sanjuán *et al*. 2004; Domingo-Calap *et al*. 2009; Peris *et al*. 2010), and a bimodal distribution with one part of the probability mass on nearly neutral mutations and the other one on the highly deleterious mutations (Hietpas *et al*. 2011). However, the data still points to a bimodal *DFE* with mutations being either neutral or very deleterious in the majority of the studies where other unimodal simpler distributions provided the best fit to the data (Sanjuán *et al*. 2004; Domingo-Calap *et al*. 2009; Peris *et al*. 2010; Jacquier *et al*. 2013). This highlights that the *DFE* might have a more complex form than the simpler probability distributions typically used to fit data. In mutation-accumulation experiments, a gamma distribution is typically assumed for the *DFE* of deleterious mutations, since there is little information to distinguish between alternative distributions (Halligan & Keightley 2009).

The other main approach is to use population genetic variation data to estimate the *DFE* with information from the site frequency spectrum (*SFS*) on putatively neutral and deleterious sites (Sawyer & Hartl 1992; Williamson *et al*. 2005; Keightley & Eyre-Walker 2007; Boyko *et al*. 2008; Gutenkunst *et al*. 2009; Kim *et al*. 2017). An interesting extension has recently been developed to take *SFS* information and divergence data from an outgroup to infer the *DFE* from the population where the *SFS* data was taken along with the rate of adaptive molecular evolution based on the divergence data (Tataru *et al*. 2017). Two other extensions have been taken to model the correlation between the fitness effects of multiple nonsynonymous alleles at a particular position (Ragsdale *et al*. 2016) and to calculate the joint *DFE* between pairs of populations (Fortier *et al*. 2019). The first step in these approaches is to inter the demographic scenario that fits the *SFS* at putatively neutral sites, which typically are chosen to be variants at synonymous sites. The *DFE* is then inferred from putatively deleterious sites of interest, typically nonsynonymous sites, while taking the demographic scenario into account. Some species where these approaches have been applied to infer the *DFE* include humans (Eyre-Walker *et al*. 2006; Boyko *et al*. 2008; Li *et al*. 2010; Huber *et al*. 2017; Kim *et al*. 2017), mouse (Kousathanas & Keightley 2013; Halligan *et al*. 2013) and *Drosophila* (Kousathanas & Keightley 2013; Huber *et al*. 2017). Studies that compare the fit of different probability distributions argue in favor of a *DFE* of deleterious nonsynonymous mutations on humans that follows either 1) a gamma distribution (Boyko *et al*. 2008; Kim *et al*. 2017) or 2) a combination of a point mass at neutrality plus a gamma distribution (Kim *et al*. 2017). Those two studies infer a leptokurtic *DFE* with a proportion of nearly neutral mutations (*s* < 10^−5^) of 18.3%-26.3%, and moderate to strong deleterious mutations (*s* > 10^−3^) of 46.6%-57.4%.

One drawback of current methods that estimate the *DFE* using population genetic variation is that they ignore all linkage information. No attempt has been made to exploit the information from linked genetic variation to estimate the *DFE* despite the fact that many studies have analyzed how both deleterious (Charlesworth *et al*. 1993, 1995; Hudson & Kaplan 1995; Nordborg *et al*. 1996; Nicolaisen & Desai 2013; Cvijović *et al*. 2018) and advantageous variants (Maynard Smith & Haigh 1974; Kaplan *et al*. 1989; Braverman *et al*. 1995; Nielsen 2005) decrease linked genetic variation. Further, linked genetic variation has been effectively used to infer the age of particular variants (Slatkin & Rannala 1997; Tishkoff *et al*. 2007; Chen & Slatkin 2013; Mathieson & McVean 2014; Chen *et al*. 2015; Nakagome *et al*. 2016; Ormond *et al*. 2016; Albers & McVean 2018), the time to the common ancestor of a positively selected allele (Smith *et al*. 2018), the time since fixation of an advantageous allele (Przeworski 2003), the selection coefficient of an allele (Slatkin 2001, 2008; Coop & Griffiths 2004; Tishkoff *et al*. 2007; Chen & Slatkin 2013; Chen *et al*. 2015; Ormond *et al*. 2016) and to detect loci under positive selection (Kim & Stephan 2002; Sabeti *et al*. 2002, 2007; Wang *et al*. 2006; Voight *et al*. 2006; Williamson *et al*. 2007; Tang *et al*. 2007; Pavlidis *et al*. 2010; Li 2011; Ferrer-Admetlla *et al*. 2014; Garud *et al*. 2015; Field *et al*. 2016; Huber *et al*. 2016). Since there has been so much success in understanding how selection changes the linked variation around individual variants, it should be feasible to pool the haplotype information from many variants putatively under selection at a certain frequency *f* to infer the distribution of fitness effects *DFE*_*f*_ of variants at a frequency *f*.

Here we propose a new approach to infer *DFE*_*f*_. We note that *DFE*_*f*_ is different from the distribution of fitness effects of new mutations entering the population, which we call the *DFE*. Natural selection acts to increase the frequency of advantageous variants and to decrease the frequency of deleterious variants, causing a difference between *DFE* and *DFE*_*f*_. The relationship between *DFE*_*f*_ and *DFE* is one of the topics we will address in this study.

Recent large population genomic datasets such as the *UK10K* (Walter *et al*. 2015), the Netherlands Genome Project (Francioli *et al*. 2014) and the Haplotype Reference Consortium (McCarthy *et al*. 2016) provide an unprecedented source of haplotype information to quantify both the *DFE*_*f*_ and the *DFE*. These datasets have started to be exploited to understand the impact of selection on variants under selection at a certain frequency. For example, Kiezun et al. (2013) found that, conditioning on the variants having a certain frequency *f* in the population, nonsynonymous variants have more extended linkage disequilibrium with neighboring neutral variation compared to synonymous variants on data from the Netherlands Genome Project. This is in line with Takeo Maruyama’s results showing that deleterious variants at a certain frequency have a younger age compared to neutral variants (Maruyama 1974), implying that there is less variation on haplotypes carrying deleterious variants.

Building on previous work to estimate the strength of selection acting on advantageous variants (Slatkin 2001; Chen & Slatkin 2013), we propose an approach to provide a point estimate of the population-scaled selection coefficient or a distribution of fitness effects acting on a set of variants *at a particular frequency f* (*DFE*_*f*_). We infer the strength of natural selection using pairwise haplotypic identity-by-state lengths (the length in one direction along a pair of haplotypes carrying a focal allele to the first difference between the pair of haplotypes). For each pair *j* of haplotypes we define the observed length as *L*_*j*_. The length can be measured in both directions along the chromosome extending outward from the focal allele. We show that these lengths can be used to distinguish between alleles under positive and negative selection in several non-equilibrium demographic scenarios. Further, we show how the *DFE*_*f*_ can be used to infer the *DFE*. The resulting method can help improve the understanding of how selection is influencing, for instance, the low-frequency variants present in a population. We apply our method to the *UK10K* dataset, and we estimate a similar proportion of neutral, moderately deleterious and deleterious variants compared to *SFS*-based approaches.

## Results

### A method for inference of the population-scaled selection coefficient based on haplotype variation

Our analysis is based on a set of *X* haplotype pairs carrying a derived allele at a frequency *f* in the population. We compute the pairwise identity by state length *L*_*j*_ for every haplotype pair, which is defined as the distance from the derived allele to the first difference between a pair of haplotypes. For computational simplicity, we bin the chromosome under analysis into a set of *S* discrete non-overlapping windows ***W*** = {*w*_1_, *w*_2_, …, *w*_*s*_} that extend to the side of the derived allele. Thus, for a set of *n* haplotype pairs carrying an allele, our analysis is based on which window the first difference appears in for each pair 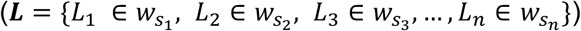. We define *s*_1_, …, *s*_*n*_ as integers between 1 and *S* indicating the windows in which each length falls (Figure 1). We can calculate a length *L*_*j*_ both upstream and downstream of each derived allele in a sample of *n* allele carriers from alleles at a frequency *f* in a number *A* of loci, and observe a total number 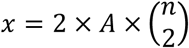 of ***L*** length values.

**Figure 1.**
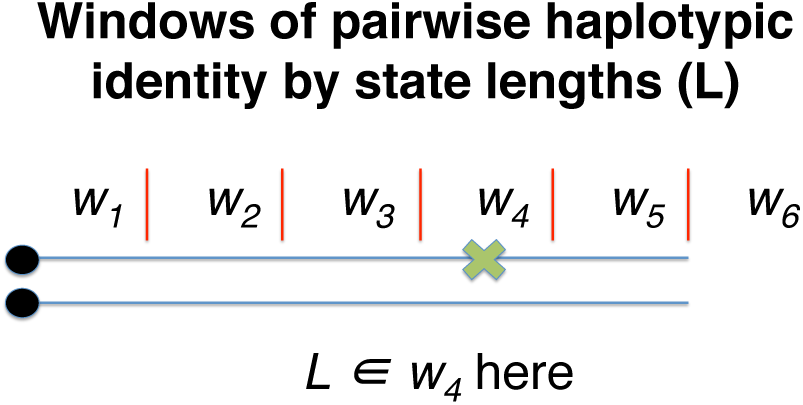
Two haplotypes containing a derived allele, here represented as a black dot, that has a frequency *f* in the population. The physical distance near the allele is divided into 5 non-overlapping equidistant windows of a certain length, with an extra window *w*_*6*_ indicating that there are no differences in any of the windows *w*_*1*_ to *w*_*5*_. The first difference between the pairs of haplotypes is denoted by the green “x”.

For our inference procedure, we will consider each *L*_*i*_ independently and so we momentarily refer generically to a single observed length as *L*. The parameter we wish to infer is the population scaled selection coefficient 4*Ns*. That parameter is defined in terms of the effective population size *N* from the most ancient epoch in the demographic scenario *D*. It is also possible to define the population scaled selection coefficient in terms of the most recent epoch. If the population size of the most recent epoch is *N*_*R*_, then the population scaled selection coefficient in the most recent time is equal to 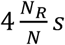.

The likelihood of a particular population scaled selection coefficient, 4*Ns*, conditioned on the allele frequency *f* and a certain demographic scenario *D*, from a single observed length *L* can be expressed as:

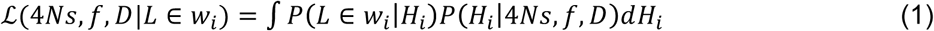

where *H*_*i*_ is a particular allele frequency trajectory. The integration over the space of allele frequency trajectories *H*_*i*_ is challenging. One possible approach to do the integration over the space of *H*_*i*_ is to perform forward-in-time simulations of alleles under the Poisson Random Field model and retain the trajectories of alleles that end at a frequency *f* in the present. However, this approach is ineffective because we will end up simulating the trajectories of many alleles that do not end up at a frequency *f* in the present. To overcome this, we integrate over the space of allele frequency trajectories *H*_*i*_ using an importance sampling approach. We also compute *P L* ∈ *w*_*i*_|*H*_*i*_ using a Monte Carlo approximation (see Methods).

We then apply this likelihood function to the complete collection of observed lengths ***L*** to calculate a composite likelihood function for 4*Ns*:

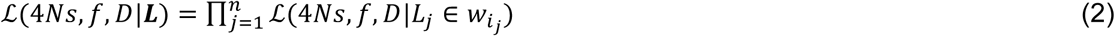

An estimator of 4*Ns* can be obtained by maximizing this composite likelihood function, which here we do simply by using a grid search over a range of candidate values (see Methods).

To build an understanding of the inference problem and the method’s performance, we first assessed the impact of selection on allele frequency trajectories, pairwise coalescent times, and haplotype identity-by-state-lengths, and then assessed the performance of the estimator. We do this first for a constant-size demographic history and then time-varying population sizes.

### Evaluation of population-scaled selection coefficient inference for constant population sizes

We investigated performance using forward-in-time simulations under the Poisson Random Field (*PRF*) framework. Specifically, we used *PReFerSim* (Ortega-Del Vecchyo *et al*. 2016) to obtain 10,000 alleles frequency trajectories with a present-day sample allele frequency of 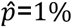 for 5 different values of selection (4*Ns* = 0, −50, −100, 50, 100) in a sample of 4,000 chromosomes (see Methods).

Using the 10,000 recorded allele frequency trajectories for each selection value *4Ns*, we calculated the mean allele frequency across many generations going backwards into the past to obtain an average frequency trajectory for 1% frequency alleles (Figure 2A). As expected, the average allele frequency trajectory for neutral alleles (4*Ns* = 0) is higher for a longer duration going backwards in time compared to alleles under natural selection. Alleles under the same absolute strength of selection have the same average allele frequency trajectory, regardless of whether the allele is under positive or negative selection. The distribution of ages is shifted towards younger values for higher absolute values of 4*Ns* and with increasingly smaller standard deviation (Figure 2B), and Maruyama’s theoretical results accurately predict the mean age estimates observed in the simulations (Supplementary Table 1).

**Figure 2.**
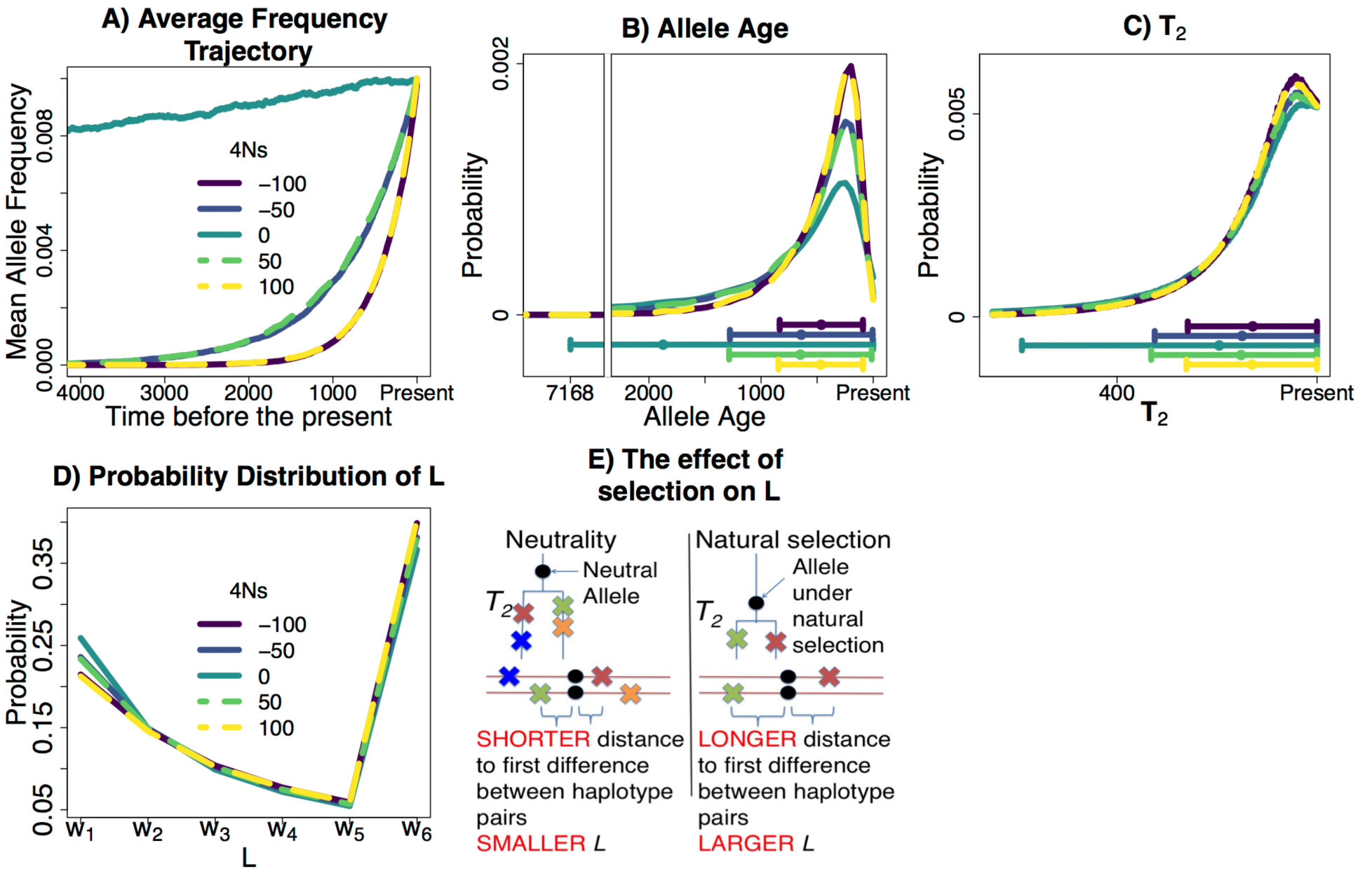
Properties of alleles sampled at a 1% frequency under different strengths of natural selection in a constant size population (*N* = 10, 000). We obtained 10,000 frequency trajectories for 1% frequency alleles under different strengths of selection using forward-in-time simulations under the *PRF* model. We used those frequency trajectories to calculate: A) The mean allele frequency at different times in the past, in units of generations, to obtain an average frequency trajectory; B) The probability distribution of allele ages; C) The probability distribution of pairwise coalescent times *T*_*2*_. Below B) and C), we show a dot with two whiskers extending at both sides of the dot. The dot represents the mean value of the distribution and the two whiskers extend one s.d. below or above the mean. The whisker that extends one s.d. below the mean is constrained to extend until max(mean – s.d., 0). D) Probability distribution of *L*. We define *L* by taking the physical distance in basepairs next to the allele across 5 non-overlapping equidistant windows of 50 kb, with an extra window *w*_*6*_ indicating that there are no differences in the 250 kb next to the allele. In this demographic scenario, the alleles under a higher absolute strength of selection have younger ages and younger *T*_*2*_ on average. The fact that alleles under higher strengths of selection have younger average *T*_*2*_ values implies that those alleles tend to have larger *L* values as shown in D) and E).

We computed the distribution of pairwise coalescent times *T*_2_ analytically (see Supplementary Methods) across different values of 4*Ns*. We found that alleles under higher absolute values of 4*Ns* have a more recent average value of *T*_*2*_, and their distribution of *T*_*2*_ has a smaller standard deviation (Figure 2C). We calculated the distribution of *L* for each 4*Ns* value using simulations assuming a constant recombination rate *ρ* = 4*Nr* = 100 and a constant mutation rate *θ* = 4*Nu* = 100 for a region of 250 kb. Alleles under the same absolute strength of selection have almost identical distributions of *L* (Figure 2D). This is in line with the fact that *T*_*2*_ is younger in alleles under stronger selection coefficients, implying that there will be fewer mutations between haplotypes sharing the allele and, therefore, higher average values of *L* (Figure 2E).

We next used the simulations to test our method’s ability to estimate the strength of selection. We found that for alleles where, for instance *4Ns* is −50, the estimated values of selection tend to be equally distributed around values of −50 or 50 (Figure 3A). A similar result is seen for the *4Ns* values equal to 100. This reinforces that in a constant size population one can only provide reasonable estimates of the absolute strength of natural selection. Indeed, when we display the estimated absolute value of the strength of selection, we see that our method produces nearly unbiased estimates (Figure 3B).

**Figure 3.**
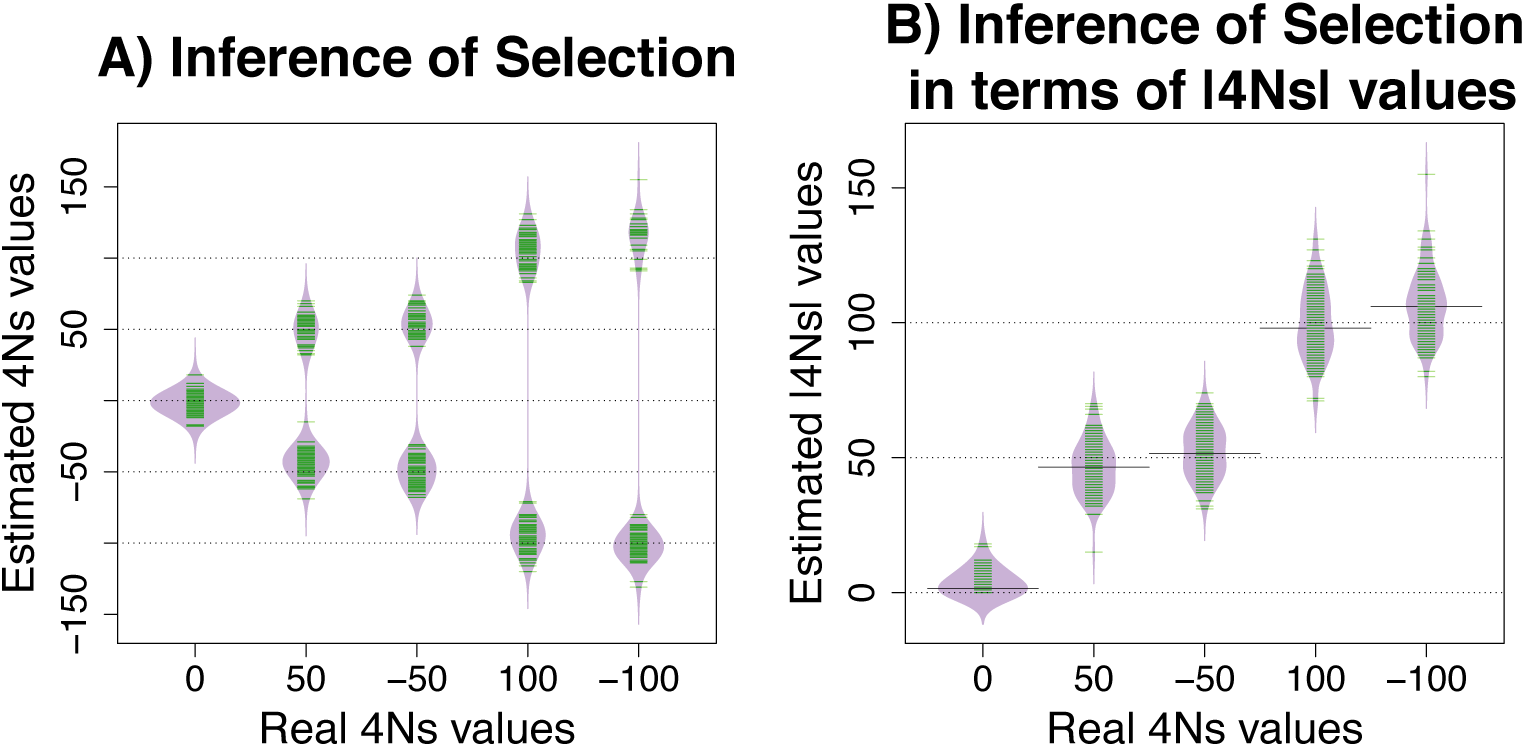
Estimation of the strength of natural selection in a constant population size model using 10, 000 realized values of *L* from 10, 000 pairs of haplotypes, where each pair was sampled from an independent loci in 1% frequency alleles. A) Estimated selection values. B) Estimated selection magnitudes (absolute values of *4Ns*). ‘Real 4*Ns* values’ refers to the 4*Ns* values used in the simulations, while ‘Estimated 4*Ns* values’ refers to the values estimated by our method. The dashed lines are placed on values that match 4*Ns* values used in the simulations. The median value of the estimates of 4*Ns* is shown with a solid line. The green lines in A) and B) indicate estimated values of 4*Ns*, where there are 100 estimated values for the five 4*Ns* values inspected. Each estimated 4*Ns* value uses 10,000 *L* values.

### Evaluation of inference performance for non-equilibrium demographic scenarios

Following our analysis for constant-size populations, we next analyzed the shape of the average allele frequency trajectory in a population expansion scenario (Figure 4A) for 1% frequency alleles with different 4*Ns* values. Unlike in the constant population size scenario, we found distinct average allele frequency trajectories for alleles under positive or negative selection (Figure 4B): alleles under positive selection on average had increased in frequency moving forward in time, while alleles under negative selection on average had increased in frequency before the expansion and then decreased after the expansion due to the increased selection efficacy in the large population. The ages of alleles under the strongest absolute values of selection tend to be younger, and alleles with the same */*4*Ns/* value but different 4*Ns* value differ in the mean and standard deviation of their allele ages (Figure 4C). The distributions of pairwise coalescent times for allele carriers show concordant patterns (Figure 4D): alleles under the stronger positive selection had, on average, younger *T*_*2*_ values than negatively selected alleles of the same magnitude. Further, when we contrasted the *T*_*2*_ distribution of the negatively selected alleles inspected (4*Ns* = −50, −100), we saw that their mean *T*_*2*_ value did not differ much, and their biggest difference was due to a slightly smaller standard deviation in the most deleterious allele (Figure 4D).

**Figure 4.**
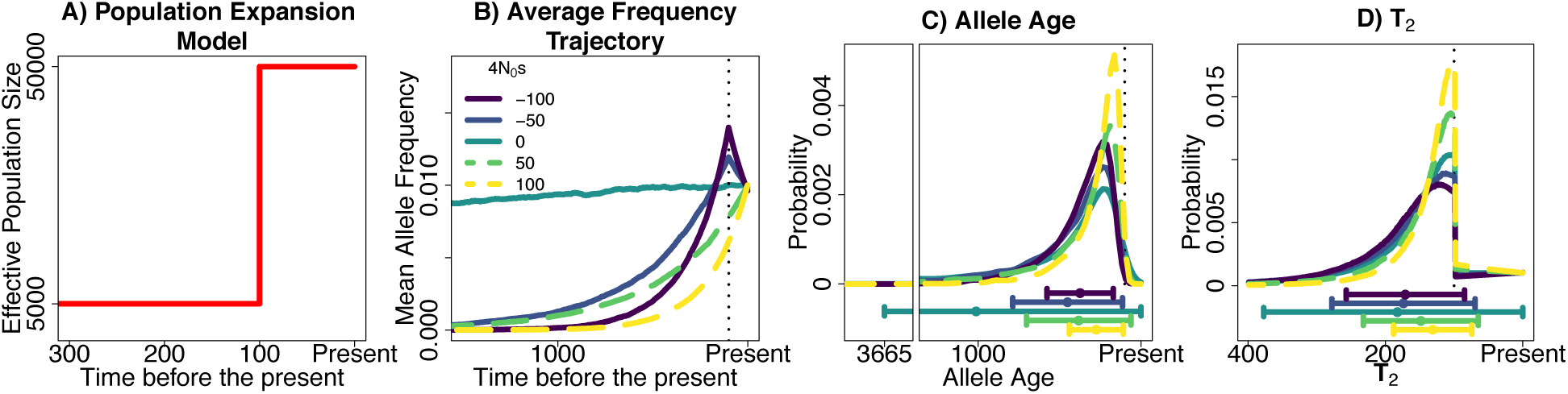
Properties of alleles sampled at a 1% frequency under different strengths of selection in a population expansion scenario. A) Population expansion model analyzed. B) Mean allele frequency at different times in the past, in units of generations. Note that alleles under the same absolute strength of selection (4*Ns*) have very different average allele frequency trajectories, in contrast to the constant population size scenario (Fig 2); C) Probability distribution of allele ages and D) Probability distribution of pairwise coalescent times *T*_*2*_. The dot and whiskers below C) and D) represent the mean value of the distribution and the two whiskers extend at both sides of the mean until max(mean +- s.d., 0).

We next used our method to infer the strength of selection for this expansion scenario and found that it can provide approximately unbiased estimates of the sign and strength of selection (Figure 5, using 10,000 realized values of *L* from 10,000 pairs of haplotypes at independent loci). This does not mean we can differentiate between positive and negative selection in all non-equilibrium models. The power to do so will be dependent on the parameters of the non-equilibrium demography being studied. As an example, in an ancient bottleneck scenario we find there are no significant differences in the distribution of *T*_*2*_ between alleles that have the same absolute strength of selection, indicating that we would not be able to differentiate between alleles under positive or negative selection under this demographic model (Supplementary Figure S1).

**Figure 5.**
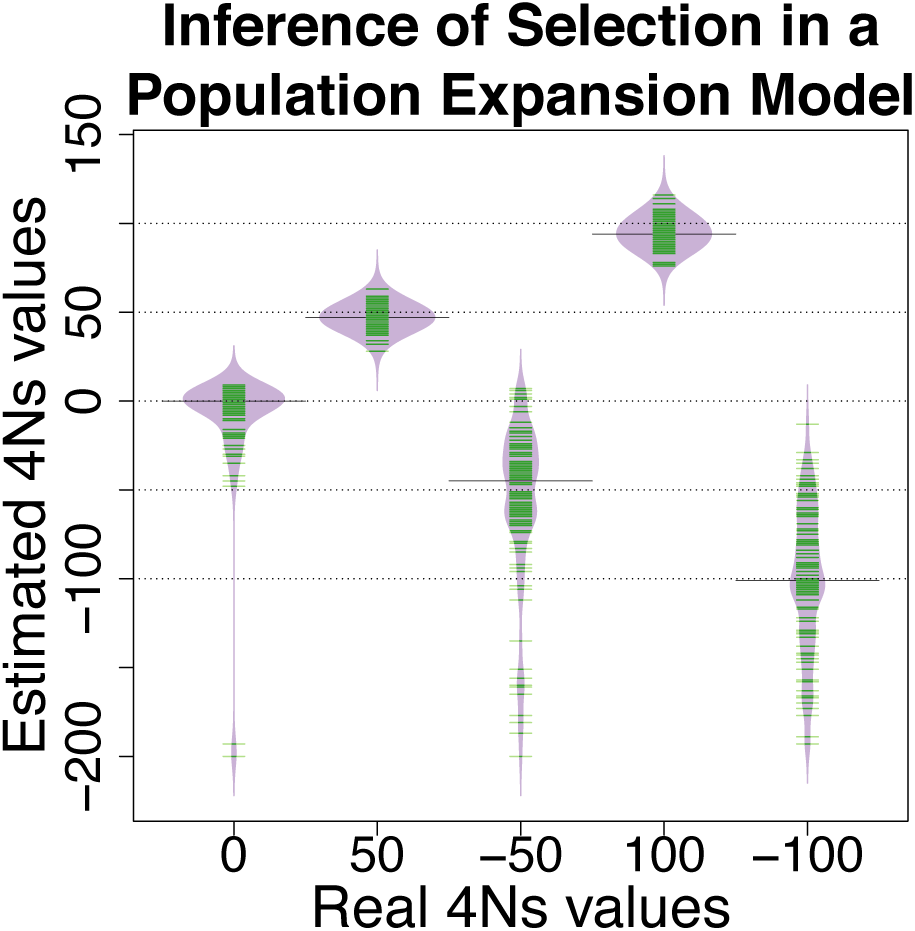
Estimation of the strength of natural selection in a population expansion model for 1% frequency alleles. The green lines indicate one estimated value of 4*Ns*. ‘Real 4*Ns* values’ indicate the 4*Ns* values used in the simulations and ‘Estimated 4*Ns* values’ refers to the values estimated by our method. The median value of the estimates of 4*Ns* is shown with a solid line. The recombination rate in the simulated 250 kb region for the most recent epoch was set equal to *ρ* = 4*Nr* = 1,000 and the mutation rate was set equal to *θ* = 4*Nu* = 1,000.

### A method for inference of the distribution of fitness effects for variants found at a particular frequency (“*DFE*_*f*_”)

Our composite likelihood framework is extendible to find the distribution of fitness effects *DFE*_*f*_ for a set of variants at a particular frequency *f*. This distribution, which we denote as *DFE*_*f*_, is different from the canonical *DFE*, which represents the distribution of fitness effects of new mutations that recently entered the population. To parameterize the *DFE*_*f*_ we use a discretized, partially collapsed gamma distribution following studies that use a gamma distribution (Boyko *et al*. 2008; Kim *et al*. 2017). We parameterize the gamma component with two parameters that represent the shape *α* and scale *β*. We discretize the distribution to cover only integer values of 4*Ns* for computational reasons, and then collapse the probabilities for all values greater than a threshold 4*Ns* value (which we denote as *τ*) to a single point mass. The point mass probability is necessary to facilitate the integration over 4*Ns* values when computing ℒ(*α, β, D, f*|*L* ∈ *w*_*i*_). We denote the resulting distribution as *DFE*_*f*_(*α, β*). In practice, we explore different values of *α* and *β* while keeping the value of *τ* fixed to a large value (i.e 300), effectively representing strongly selected variants (see Methods).

The likelihood of having a certain distribution of identity by state lengths *L* given a demographic scenario *D*, a variant at a frequency *f* and two parameters *α* and *β* is equal to:

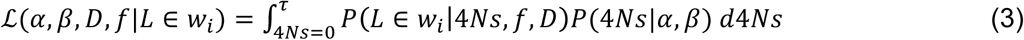

Where *P* (*L* ∈ *w*_*i*_ |4*Ns, f, D*) = ℒ(4*Ns, f, D*|*L* ∈ *w*_*i*_) and was introduced in equation 1.

### Testing the inference of the distribution of fitness effects for variants found at a particular frequency (“*DFE*_*f*_”)

We tested if the distribution of haplotype lengths *L* can be used to estimate the parameters that define the distribution of fitness effects of variants at a particular frequency. We used distributions of 100,000 *L* values obtained via simulations under the constant population size and population expansion demographic model from the past sections under two distributions of fitness effect of new mutations estimated in different species: one from humans (shape = 0.184; scale = 319.8626; N = 1000) (Boyko *et al*. 2008) and another one from mice (shape = 0.11; scale = 8636364; N = 1000000) (Halligan *et al*. 2013).

We found that the estimated parameters of the shape (*α*) and scale (*β*) of the *DFE*_*f*_ of 1% frequency variants in a sample of 4,000 chromosomes have considerable variation (Figure 6A,B). However, the estimated shape and scale of the *DFE*_*f*_ typically imply the correct mean value of the *DFE*_*f*_ (estimates lie along the red-dashed lines in Figure 6). This can be better seen in Supplementary Figure S2. We found that the estimated *DFE*_*f*_ parameters on constant population sizes define a *DFE*_*f*_ with a mean 4*Ns* value that, on average, is almost equal to the mean 4*Ns* value found across 50,000 simulated 1% frequency variants. In a population expansion scenario (Figure 6C,D), the estimated *DFE*_*f*_ parameters imply a *DFE*_*f*_ with a mean 4*Ns* value that is slightly lower than the actual mean 4*Ns* value, and with considerably higher variance in the estimated mean (Supplementary Figure S2).

**Figure 6.**
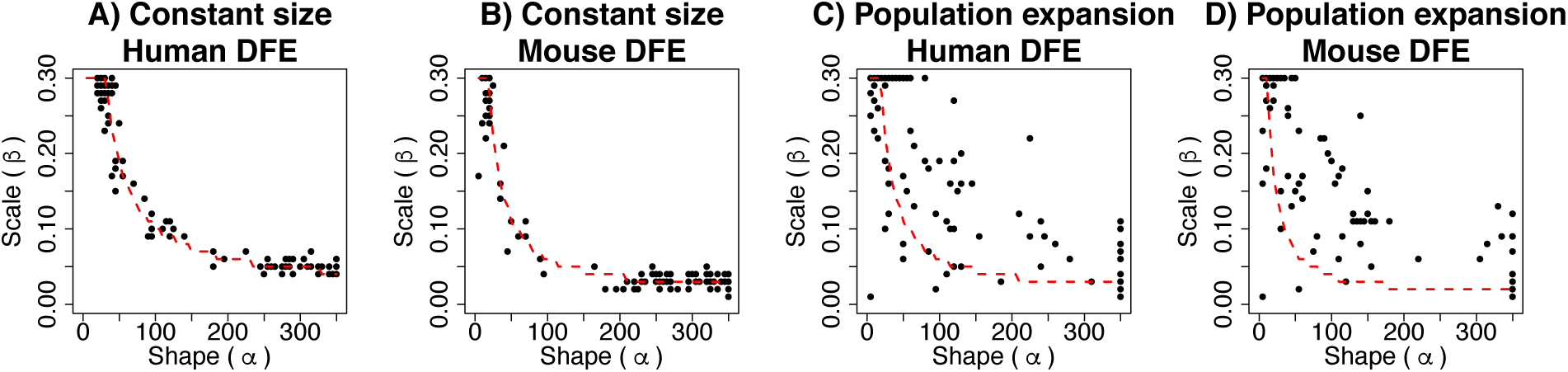
MLEs of the parameters that define the distribution of fitness effect for variants at a 1% frequency. We tested if our method was capable of estimating the parameters of the *DFE*_*f*_ of variants at a particular frequency in two demographic models and two *DFE*’*s*. The shape (*α*) and scale (*β*) parameters define the compound *DFE*_*f*_ distribution. Each black dot represents the *α* and *β* parameter estimated using a set of 100,000 *L* values simulated independently. The dotted red line represents a combination of shape and scale parameters from a gamma distribution that give an identical mean *4Ns* value to the mean *4Ns* value of the underlying *DFE*_*f*_. The grid of scale parameters explored goes from (0.01, 0.02, …, 0.3) and the grid of shape parameters explored goes from (5, 10, …, 350).

### Method for inferring the distribution of fitness effects of new mutations (*DFE*) from the distribution of fitness effects for variants at a particular frequency (*DFE*_*f*_)

The distribution of fitness effects of variants at a particular frequency (*DFE*_*f*_) is related to the distribution of fitness effects of new variants *DFE* by equation 4 (see Methods for more detail):

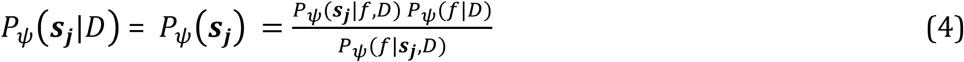

where ***s***_***j***_ is an interval of 4*Ns* values [4*Ns*_*0*_, 4*Ns*_*1*_). *s*_*0*_ and *s*_*1*_ define two different selection coefficients. We used a set of non-overlapping intervals *s = {[*4*Ns*_*0*_, 4*Ns*_*1*_*), [*4*Ns*_*1*_, 4*Ns*_*2*_*), [*4*Ns*_*2*_, 4*Ns* _*3*_*)…*, [ 4*Ns* _*b-1*_, 4*Ns* _*b*_*)} = {* ***s***_**1**_, ***s***_**2**_, ***s***_**3**_, …, ***s***_***b***_ *}. ψ* is a vector of the parameters *ψ* = {*ψ*_1_, *ψ*_2_, *ψ*_3_, …, *ψ*_k_} that define the *DFE*.

The probabilities *P*_*ψ*_(***s***_***j***_|*f, D*) over all the intervals in *s* define the distribution of fitness effects of variants at a particular frequency *DFE*_*f*_ over a set of discrete bins. After inferring the *DFE*_*f*_ using our composite likelihood method, we can calculate *P*_*ψ*_(***s***_***j***_|*f, D*) from the inferred *DFE*_*f*_. On the other hand, *P*_*ψ*_ (***s***_***j***_|*D*) = *P*_*ψ*_(***s***_***j***_) since the demographic scenario *D* does not change the proportion of new variants in a selection interval ***s***_***j***_. *P*_*ψ*_(***s***_***j***_) defines the proportion of new mutations inside a ***s***_***j***_ interval. It is equal to the *DFE* over a set of discrete intervals ***s***_***j***_. Regarding the other two probabilities shown in the equation, *P*_*ψ*_(*f*|*D*) can be estimated by measuring the proportion of variants at a certain frequency *f* given *D* and a set of parameters ***ψ*** that define the *DEF. P*_*ψ*_(*f*|***s***_***j***_, *D*) can be computed via simulations (see Supplementary Text for more details).

### Testing inference of the distribution of fitness effects of new mutations *DFE* from the distribution of fitness effects of variants at a particular frequency (*DFE*_*f*_)

We estimated the distribution of fitness effects of new mutations, i.e. the *DFE*, in a population expansion scenario given the distribution of fitness effects *DFE*_*f*_ of a set of variants at a 1% frequency (Figure 7 – Boyko Human *DFE*; and Supplementary Figure S3 – Human *DFE* with a scale value that is 20 times smaller). We see that the inferred and real *P*_*ψ*_(***s***_***j***_) values match using equation (4), with some slight discrepancies that could be due to either using a ***s***_***j***_ bin that is not small enough or small inaccuracies in the estimated probabilities of *P*_*ψ*_(***s***_***j***_|*f, D*), *P*_*ψ*_(*f*|*D*) or *P*_*ψ*_(*f*|***s***_***j***_, *D*). We also note that variants at a 1% frequency tend to be less deleterious compared to new variants based on the comparison of the distributions *P*_*ψ*_(***s***_***j***_|*f, D*) against *P*_*ψ*_ (***s***_***j***_). Additionally, we used our *DFE*_*f*_ estimates from Figure 6 to estimate *P*_*ψ*_(***s***_***j***_). The *P*_*ψ*_(***s***_***j***_) estimates are accurate, but display a larger variance under the population expansion scenario compared to the constant size scenario (Supplementary Figure S4).

**Figure 7.**
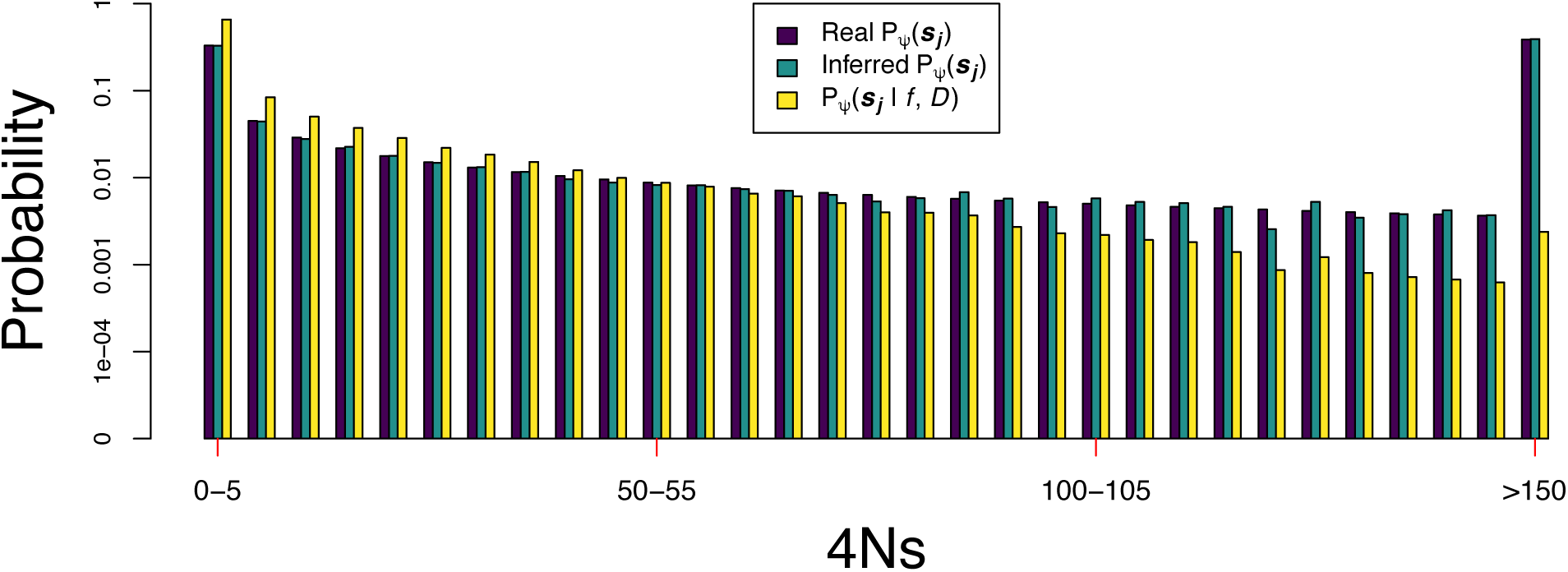
Inference of the distribution of fitness effects of new mutations from the distribution of fitness effects of variants at a certain frequency in deleterious variants. The *DFE* follows a gamma distribution with shape and scale parameters equal to 0.184 and 1599.313, respectively. This is equal to the gamma distribution inferred by Boyko et al. (2008) after adjusting the population sizes to the population expansion demographic model used. The demographic model has a population that grows from 5,000 to 50,000 individuals in the last 100 generations (see also Figure 4A). ‘Real ***P***_***ψ***_ (***s***_***j***_) *‘* refers to the probability of having a 4*Ns* value in a certain interval ***s***_***j***_ given the distribution of fitness effects of new mutations with parameters ***ψ***. ‘***P***_***ψ***_(***s***_***j***_|***f, D***)*’* is the probability of having a 4*Ns* value in an interval ***s***_***j***_ given the distribution of fitness effects *DFE* with parameters ***ψ*** and the demographic scenario *D* in *f* = 1% frequency variants. We calculated ***P***_***ψ***_(***s***_***j***_|***f, D***) from a set of ∼ 40,000 4*Ns* 1% variants obtained via *PReFerSim* simulations under the *DFE* and the population expansion scenario (see Supplementary Text). ‘Inferred ***P***_***ψ***_ (***s***_***j***_) *‘* is an estimate of the probability of having a 4*Ns* value in a certain interval ***s***_***j***_ given the distribution of fitness effects of new mutations with parameters ***ψ*** using ***P***_***ψ***_(***s***_***j***_|***f, D***) and equation 4. The selection coefficient *s* refers exclusively to the action of deleterious variants in this plot.

### Application: Inference of the distribution of fitness effects of 1% frequency variants in the UK10K dataset

We inferred the distribution of fitness effects of the 273 1% ± 0.05% frequency variants at non-CpG nonsynonymous sites that are more than 5 Mb away from the centromere or telomeres in the phased *UK10K* haplotype reference panel. The panel was statistically phased with *Shapeit2* (Delaneau *et al*. 2013b), which previous analyses have shown produces a low haplotype phasing error (switch error rate approximately < 2.0%) for low-frequency alleles (Delaneau *et al*. 2013a). Our method assumes that phasing errors will be similar in the nonsynonymous and synonymous variants, implying that differences in the distribution of *L* will be due to selection instead of phasing errors. We discarded a set of related individuals along with other individuals with no clear European ancestry from the haplotype panel, as previously defined (Walter *et al*. 2015). In the end, we obtained a set of 3,621 individuals (7,242 haplotypes) from the *UK10K* haplotype panel.

We used an *ABC* algorithm to infer the demographic scenario that explains the distribution of *L* for the 152 non-CpG synonymous variants at a 1% ± 0.05% frequency that are more than 5 Mb away from the centromere or telomeres (see Supplementary Methods, Supplementary Figure S5). CpG sites were removed before estimating *L* around the non-CpG synonymous sites. We removed CpG sites by excluding sites preceded by a C or followed by a G (McVicker *et al*. 2009). Due to computational reasons, in the *ABC* method we scaled the population size down by a factor of five while increasing the mutation rate *μ*, selection coefficient *s* and recombination rate *r* by the same factor of five to keep 4*Ns, Θ* = 4*Nμ* and *ρ* = 4*Nr* constant. That same scaling was used in all the simulations described in this section and in our inference of selection in the *UK10K* data. We will refer to the inferred scaled model as the ‘scaled *UK10K* model’ and we will refer to the model without the scaling as the ‘*UK10K* model’. We find that in the upstream and downstream 250 kb regions surrounding the 152 synonymous 1% frequency variants and the 273 nonsynonymous 1% frequency sites there is a similar proportion of exonic sites (Mann-Whitney U test p-value = 0.876), PhastCons element sites (Mann-Whitney U test p-value = 0.299), and the average strength of background selection (Mann-Whitney U test p-value = 0.605) based on the *B* values (McVicker *et al*. 2009). The distributions of *B* values indicate that similar strengths of background selection are acting on the synonymous and nonsynonymous sites, and should reduce genetic variation similarly on regions surrounding both categories of sites. Therefore, the demographic model we inferred for the synonymous variants can be used to model the evolution of the nonsynonymous variants since the reduction in genetic variation due to background selection is similar on the haplotypes surrounding both types of variants (Supplementary Figure S6). The approach of inferring the demographic model using synonymous sites is not novel for analyses with the site frequency spectrum and helps control for the effects of background selection (Boyko *et al*. 2008; Huber *et al*. 2017; Kim *et al*. 2017; Tataru *et al*. 2017).

We performed simulations under the scaled *UK10K* model inferred using the *ABC* algorithm. We found that the frequency trajectories and allele ages are significantly different between alleles under different strengths of selection (Figure 8). However, the distribution of *T*_*2*_ values is very similar for deleterious alleles that experience up to a twofold difference in the amount of selection acting upon them. This is important to note since the distribution of *T*_*2*_ values is one of the most important factors, along with the mutation and recombination rate, determining the resolution of our approach to infer selection.

**Figure 8.**
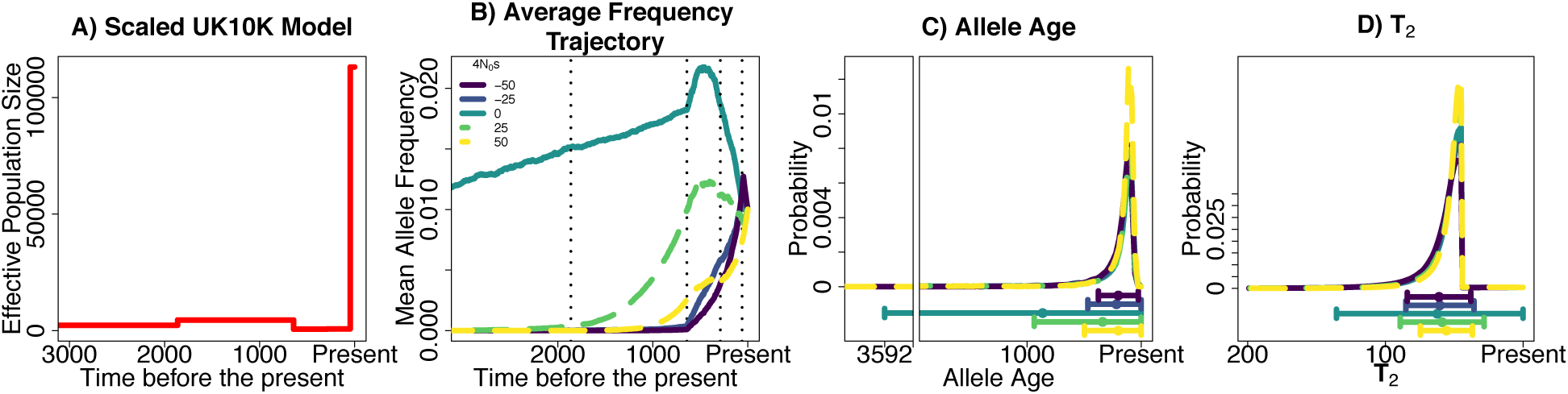
Properties of alleles sampled at a 1% frequency under different strengths of natural selection in the scaled *UK10K* model inferred in the *UK10K* data. A) Population model inferred in the *UK10K* dataset. B) Mean allele frequency at different times in the past, in units of generations. C) Probability distribution of allele ages and D) Probability distribution of pairwise coalescent times *T*_*2*_. The dot and whiskers below C) and D) represent the mean value of the distribution and the two whiskers extend at both sides of the mean until max(mean +- s.d., 0).

We also performed simulations to analyze if the amount of information present in the *UK10K* dataset was sufficient to infer selection coefficients in 1% frequency variants. Our approach takes into account the differences in recombination rates on the regions surrounding each variant on the genome in the *UK10K* data (Supplementary Methods). We performed 100 simulation replicates, where each replicate mimics the amount of information present in the *UK10K* dataset. Each replicate contains 273 independent loci with 72 haplotypes containing the derived allele. The recombination rates, both to the left and right side of the loci, were assigned based on the average per base recombination rate in the 250 kb region surrounding each variant (see Supplementary Figure S7). We calculated *L* moving upstream and downstream of the focal loci, obtaining 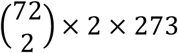 values for each simulation replicate. Using data simulated under 5 different selection coefficients, we found that we were able to obtain accurate estimates of selection when the variants were neutral or under positive selection. When we simulated deleterious variants, we found that our estimates of selection tended to be biased towards being more neutral than the actual 4*Ns* value. However, the true value was within the 10^th^ and 90^th^ percentile of the distribution of estimated values (Supplementary Figure S8). We obtained similar results when the simulated 273 loci shared the same recombination rate (Supplementary Figure S9). We obtained equally accurate estimates of *P*_*ψ*_(***s***_***j***_) on the ***s***_***j***_ intervals when we performed simulations using the Boyko distribution of fitness effects under the scaled and *UK10K* demographic model (Supplementary Figure S10-S11; Supplementary Table S2-S3).

We performed bootstrap replicates of the *L* values from the 273 1% frequency nonsynonymous variants of the *UK10K* dataset and the 152 1% frequency synonymous variants to evaluate the variation in our estimates of 4*Ns*. We removed CpG sites before estimating the *L* values surrounding the nonsynonymous and synonymous variants. The variation around the estimates using bootstrap replicates is shown in Supplementary Figure S12, where we see that the point estimates in the replicates tend to be close to a 4*Ns* value equal to 0 for both nonsynonymous and synonymous variants. We performed the inference on the 1% frequency synonymous variants because an inferred 4*Ns* value that was nominally different from 0 would indicate problems with our methodology such as a misspecified demographic model.

We used the *L* values for the 273 nonsynonymous variants at a 1% frequency to infer the parameters of the distribution of fitness effects *DFE*_*f*_. We assume that no derived variants we observe are under positive selection and that the *DFE*_*f*_ follows a gamma distribution with a point mass, as explained in the section *Inference of the distribution of fitness effects of variants at a particular frequency*. When we solved the integral from Equation 3, we used discretized values of 4*Ns* that went from 0 to 75, and we defined that 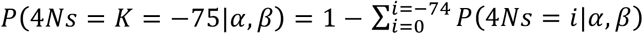. We only explored 4*Ns* values from 0 to −75 because we only had high resolution for those 4*Ns* values (as indicated by *ESS* values bigger than 100, see Supplementary methods for an explanation of *ESS* values; Supplementary Figure S13). We inferred a scale value of 0.01 and a shape value of 0.03. Based on a set of bootstrap replicates, we found that our estimates clustered on the edges of the shape parameter values explored (Supplementary Figure S14). This effect is specific to the inferred demographic scenario for the *UK10K* dataset, since we did not observe the same phenomenon in the simulations done under the constant population size and population expansion demographic scenarios we explored previously (Figure 6). Based on our estimates of the *DFE*_*f*_, we estimated *P*_*ψ*_(***s***_***j***_) by employing Equation 4 and using *P*_*ψ*_(*f*|*D*) (see Supplementary Methods for an explanation of our calculation of *P*_*ψ*_(*f*|*D*)). We compared those values with previously obtained estimates (Boyko *et al*. 2008; Kim *et al*. 2017). The point estimates of *P*_*ψ*_(***s***_***j***_) along with the 90% bootstrap percentile intervals for other ***s***_***j***_ intervals are shown in Figure 9 and Supplementary Figure S15. We also show information for other bootstrap percentile intervals on Supplementary Table S4. Based on our 90% bootstrap percentile intervals we find that our estimate of *P*_*ψ*_(***s***_***j***_ ∈ [5,50)) is smaller than the probabilities computed by Boyko *et al*. 2008 and Kim *et al*. 2017. On the other hand, the estimate of *P*_*ψ*_(***s***_***j***_ ∈ [50, ∞)) was bigger than the estimates of Boyko *et al*. 2008 and Kim *et al*. 2017. The probabilities of having a value of selection *s* over different orders of magnitude are shown on Supplementary Table S5 and are compared with the probabilities obtained by (Boyko *et al*. 2008; Kim *et al*. 2017). We also computed p-values under the null hypothesis that there is no difference between the estimated *P*_*ψ*_(***s***_***j***_) values from the data and the *P*_*ψ*_(***s***_***j***_) from the Boyko distribution of fitness effects (see Supplementary Figure S16). The p-values were bigger than 0.05 for the three intervals ***s***_***j***_ ∈ [0,5), ***s***_***j***_ ∈ [5,50) and ***s***_***j***_ ∈ [50, ∞). Therefore, the distribution of fitness effects is not different from the distribution of fitness effects estimated by Boyko et al. (2008) over the three ***s***_***j***_ intervals inspected.

**Figure 9.**
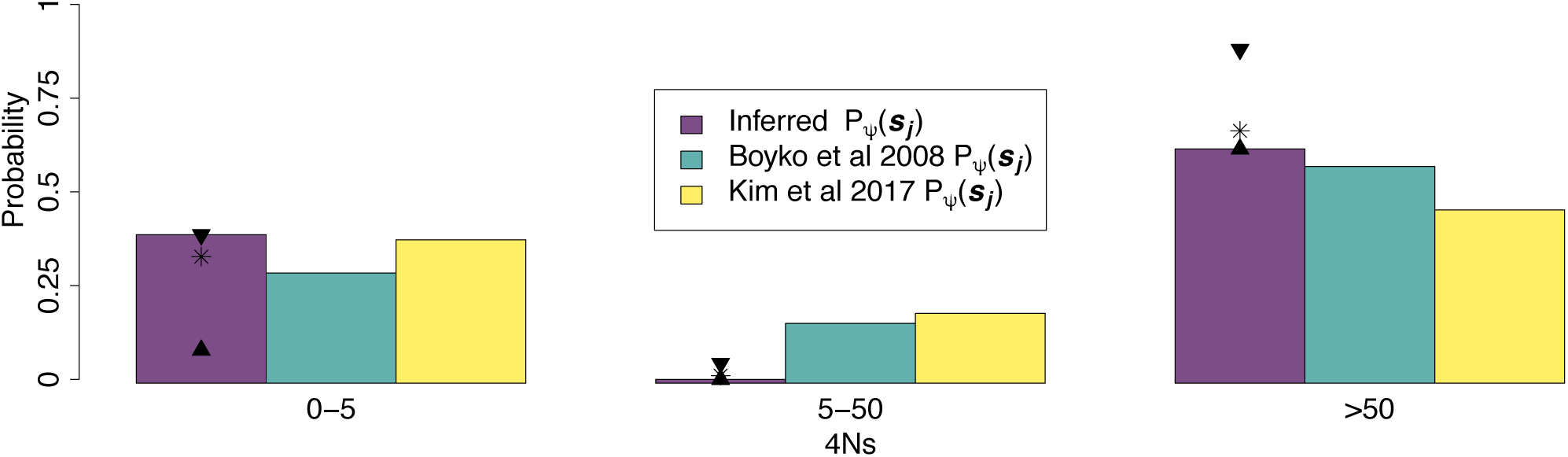
Inferred distribution of fitness effects of new mutations and 1% frequency deleterious variants in the *UK10K* dataset. ‘Inferred *P*_***ψ***_(***s***_***j***_) *‘* refers to the probability of having a 4*Ns* value in a particular interval ***s***_***j***_ given the distribution of fitness effects of new mutations *DFE*. We estimated *P*_***ψ***_(***s***_***j***_) for the ***s***_***j***_ interval = [5, 50) by summing up the *P*_***ψ***_(***s***_***j***_) probabilities over the invervals [5, 10), [10, 15), [15, 20), [20, 25), [25, 30), [30, 35), [35, 40), [40, 45) and [45, 0). The selection coefficient *s* refers exclusively to the action of deleterious variants in this plot. We compared our inferences with those of Boyko et al. (2008) and Kim et al. (2017). The two triangles shown in each ***s***_***j***_ interval denote the upper and lower limit of the 90% bootstrap percentile interval across 100 bootstrap replicates. The asterisk signs are the mean values for the inferred probabilities *P*_***ψ***_(***s***_***j***_) calculated from 100 bootstrap replicates. Despite the fact that the estimated Boyko et al 2008 *P*_***ψ***_(***s***_***j***_) values fall outside of the 90% bootstrap percentile from the inferred *P*_*ψ*_(***s***_***j***_) in the intervals ***s***_***j***_ ∈ [5,50) and ***s***_***j***_ ∈ [50, ∞), these differences are not significant according to p-values computed under the null hypothesis that there is no difference between the estimated *P*_*ψ*_(***s***_***j***_) values and the *P*_*ψ*_(***s***_***j***_) from the Boyko distribution of fitness effects (see Supplementary Figure S16).

## Discussion

We have developed a composite likelihood method to estimate the strength of natural selection acting on alleles at a certain frequency in the population. Our method builds upon previous work showing signatures of higher linkage disequilibrium for putatively deleterious alleles in comparison with neutral alleles (Kiezun *et al*. 2013). This result was shown to be in line with Takeo Maruyama’s work showing that deleterious alleles at a certain frequency tended to be younger than neutral alleles in constant population sizes (Maruyama 1974). Here we introduce a method to estimate the strength of natural selection based on linkage disequilibrium using the pairwise identity by state lengths *L*.

We found that the distribution of *L* captures differences in the absolute strength of the selection coefficient 4*Ns* in a constant population size scenario. The mean allele frequency trajectory is practically identical for deleterious and advantageous alleles experiencing the same amount of selection; therefore, any statistic based on haplotype signatures will be insufficient in that scenario to distinguish between positive and negative selection.

On the other hand, we found that the distribution of *L* is sufficient to differentiate between advantageous and deleterious alleles under some non-equilibrium demographic scenarios, including the demographic scenario inferred from the *UK10K* dataset. This is encouraging, since most natural populations are very likely to have evolved under a non-equilibrium demographic scenario and it is precisely in such scenarios where we would like to be able to differentiate between alleles with different types of selection.

The mean allele frequency trajectories of deleterious alleles segregating at a 1% frequency when the population is expanding are particularly noteworthy. These alleles tend to have increased in frequency when the population size is low. Then, they decrease in frequency when the population expands due to a higher efficacy of selection. This suggest that it is likely that, on average, deleterious alleles would tend to come from higher frequencies in the recent past in expanding populations. These simulations of allele frequency trajectories under several demographic scenarios are useful to understand past fluctuations in frequency and haplotypic patterns one might expect for selected alleles. Recent work has analyzed how different summaries of genetic variation change over time in non-equilibrium scenarios (Peischl *et al*. 2013; Lohmueller 2014a; Simons *et al*. 2014; Do *et al*. 2015; Henn *et al*. 2015; Balick *et al*. 2015; Brandvain & Wright 2016; Marsden *et al*. 2016; Koch & Novembre 2017), and analyzing the behavior of frequency trajectories is helpful to understand those changes.

When we estimated parameters that define the *DFE*_*f*_ of segregating variants, we found that our method can provide reasonable estimates of the parameters that would lead to estimating a sensible value of the mean of the *DFE*_*f*_ in several scenarios. Under a constant population size, the scale estimates of the *DFE*_*f*_ are inversely correlated with the shape parameters. Note that this curve decay causes the product of the scale and shape parameters to have relatively similar values. Under a population expansion model, the estimates of the shape and scale show a wider variation around the curve than the constant population size scenario (Figure 6). Similarly, the pairwise coalescent time *T*_*2*_ distribution between variants with different negative selection coefficients appear more similar to each other in a population expansion scenario as compared to a constant population size scenario (Figure 4D and 2C). Due to the greater variation in the estimates of the parameters that define the *DFE*_*f*_ of variants at a 1% frequency, we also see a larger variation in the mean 4*Ns* values estimated in a population expansion as compared to a constant population size demographic scenario (Supplementary Figure S2). Estimates of the mean 4*Ns* values are more precise under a constant population size compared to the population expansion scenario. For the *UK10K* demographic scenario and the scaled *UK10K* model, where there is a large recent population expansion, we saw that the proportion of 4*Ns* values smaller than 5 tended to be overestimated while the proportion of 4*Ns* values larger than 5 were underestimated based on the analysis of simulations using the Boyko et al. (2008) *DFE*. The consequence is that the mean 4*Ns* value would tend to be underestimated under the *UK10K* demographic scenario and the scaled *UK10K* demographic scenario (Supplementary Figure S10-S11). It is likely that this underestimation will be seen in other scenarios with large recent population expansions.

One technical aspect from our methodology that could be subject to future improvement is that the space of scale and shape parameters we explore is limited due to low effective sample size (*ESS*) values. In the case of the UK10K dataset, the *ESS* are smaller than 100 in 4*Ns* values smaller than −75 (Supplementary Figure S12). To increase the values of the *ESS*, one possible improvement of our method is to make better proposals for the allele frequency trajectories going backwards in time. That is, to improve our choice of the importance sampling distribution. Future work will be devoted to make improvements in this issue, particularly in populations undergoing recent large expansions. One possibility is to expand the theory of Wright-Fisher bridges to select trajectories that end at a certain frequency *f* in the present under non-equilibrium scenarios (Schraiber *et al*. 2013). We did not find the same pattern of low *ESS* values in the other two demographic scenarios we analyzed, where the population sizes did not experience changes in population size of the same magnitude as in the demographic model inferred in the *UK10K* data.

Using the *UK10K* data, we obtained a point estimate, along with 90% bootstrap interval calculations, of the *DFE*. Our point estimates are consistent with point estimates obtained using information from the site frequency spectrum (Boyko *et al*. 2008) (Supplementary Figure S16). It is possible that we find discrepancies between the estimated *DFE* in other species or populations using haplotypic information compared to using data from the site frequency spectrum. In a similar vein, important discrepancies on the inferred past demographic histories on human populations have been found when using site frequency spectrum data and haplotypic information, and some of the potential causes of the differences have been carefully discussed previously (Harris & Nielsen 2013; Hsieh *et al*. 2016; Beichman *et al*. 2017). Technical aspects of the data that can impact the demographic inferences when using haplotypic data include: 1) Switch errors during statistical phasing which cause a bias towards more recent split-time estimates (Song *et al*. 2017), 2) Uncalled heterozygous sites due to low genomic coverage which causes a bias towards lower effective population size estimates (Nadachowska-Brzyska *et al*. 2016), 3) Not filtering low coverage, potentially false positive variants, which can produce poor estimates of sudden contractions or expansions (Nadachowska-Brzyska *et al*. 2016).

With respect to the potential impact of switch errors in our inference, the *UK10K* project does not report switch error rates, but we would expect them to be even lower than those of the 1000 Genomes Project (estimated to be 0.56% with a mean of distance of ∼1,062 kb between errors) (Auton *et al*. 2015), due to the fact that the *UK10K* has approximately 50% more samples than the 1000 genomes project, and all the samples come from the same population. We expect to see the impact of phasing errors to be small in our data since we are using window sizes of 500 kb in our analysis; this window size is smaller than the mean distance between switch errors in the 1000 Genomes Data, and the mean distance between switch errors is likely to be even larger in the *UK10K* project.

Our inferences of the *DFE* can be impacted due to the low genomic coverage present in the *UK10K* dataset **(∼4x** on average). However, the estimate of the percentage of genotypes correctly called in the *UK10K* dataset is equal to 99.688% for common variants with a frequency bigger than 5%, and 99.999% for singletons (Walter *et al*. 2015). This indicates that the sequencing strategy carried out in the *UK10K* dataset should not have a large impact on our estimates of the *DFE* due to wrongly called genotypes across individuals.

Apart from the technical aspects that could be impacting our estimates of the *DFE*, there are biological phenomena that could be responsible for differences in the *DFE* estimates we see when we use site frequency spectrum information and haplotypic data. One of those phenomena is linked selection, which reduces the genetic variation in neutral sites next to an allele under either positive or negative selection (Cutter & Payseur 2013). Linked selection will increase the lengths of the pairwise haplotype lengths in the synonymous sites used to infer the demographic scenario and in the nonsynonymous sites used to infer the distribution of fitness effects. Previous work estimating the distribution of fitness effects using site frequency spectrum information has shown that using synonymous sites to estimate the demographic scenario controls for the effect of linked selection and gives an accurate estimation of the *DFE* (Huber *et al*. 2017). We expect the same effect to take place when using haplotypic information. Specifically the amount of linked selection is predicted to be similar between synonymous and nonsynonymous variants at 1% frequency (see caption Supplementary Figure S6), indicating that the increase in pairwise haplotype lengths should be similar for both synonymous and nonsynonymous sites.

Another biological phenomenon that could impact our *DFE* estimates is the incompleteness of the demographic model fitted to the data (Harris & Nielsen 2013; Garud *et al*. 2015; Beichman *et al*. 2017). We are fitting a demographic model with one deme to the *UK10K* dataset, and it is possible that fitting a model with population structure could give a better fit to the haplotypic data and to the site frequency spectrum data (Harris & Nielsen 2013). We also are not modelling non-crossover gene conversion (Andolfatto & Nordborg 1998; Korunes & Noor 2017). Non-crossover gene conversion events involve haplotype tracts of approximately 100-1000 bp and the probability that any site in the genome is involved in a non-crossover gene conversion event is 5.9 *×* 10^−6^ / bp / generation (Williams *et al*. 2015). Their impact is to break down linkage disequilibrium, which in our model, for a single variant would result in inferences that are biased towards neutrality; however, in aggregate if it impacts LD around synonymous and nonsynonymous variants equally, the effect on inferences may be minor. Nonetheless, modelling noncrossover gene conversion could improve models of the haplotype signatures of selection.

As another factor, changes on the *DFE* over time could lead to differences in the inferred *DFE* from the site frequency spectrum and the haplotypic data. *DFE* estimates from the site frequency spectrum data use information from variants that have appeared across a broad range of time. On the other hand, the haplotype data we used comes from 1% frequency variants that have appeared recently. The relaxation of selective pressures across time is one way to change the selective coefficient of variants to make them more neutral (Somel *et al*. 2013; Lynch 2016). Our results argue in favor of conserved selective coefficients over time in humans, in line with recent results (Fortier *et al*. 2019).

Although here we analyzed the distribution of fitness effects of nonsynonymous variants at a certain frequency, it is possible to determine the distribution of fitness effects of variants within specific functional categories. One possibility is to try to determine the strength of selection of alleles on variants that are predicted to be more deleterious based on the Fitcons (Gulko *et al*. 2015), SIFT (Sim *et al*. 2012), Polyphen (Adzhubei *et al*. 2010) or C-scores (Kircher *et al*. 2014; Racimo & Schraiber 2014). It is also be possible to estimate the strength of selection in a set of alleles that have a particular collection of genomic features (Huang & Siepel 2019). This can help us to obtain genome-wide estimates of the selection coefficient of variants based on their predicted functional category. This is of particular interest to genome-wide association studies, due to the interest in understanding the association between associated variants and their selection coefficients on different complex traits. Additionally the use of the newly developed tree-sequence framework (Kelleher *et al*. 2018; Haller *et al*. 2019) for simulations should also help to speed up the calculation of the likelihood of different values of selection in the part of our method that depends on Monte Carlo simulations. Another future avenue of research is to infer the distribution of selection coefficients of new mutations combining information from the *DFE*_*f*_ inferred at many different frequencies in the population. Combining information from variants at many frequencies is likely to increase the accuracy of estimates of the distribution of fitness effects of new variants, and could detect changes in the distribution of fitness effects of new variants through time.

## Methods

### Inference of selection

The likelihood of having a particular selection coefficient 4*Ns* conditioning on the allele frequency *f* and the demographic scenario *D* using information from one length *L* ∈ *w*_*i*_ can be estimated as:

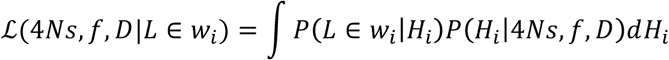

where *H*_*i*_ is a particular allele frequency history, i.e. a trajectory of allele counts from when the allele first appears in the population until the present. We can compute *P* (*L* ∈ *w*_*i*_ |*H*_*i*_) via Monte Carlo simulations done using *mssel* (Kindly provided by Richard Hudson), which assumes the structured coalescent model to simulate haplotypes containing a site whose frequency trajectory is determined by *H*_*i*_. We used *mssel* to simulate many pairs of haplotypes (10,000 independent pairs for all scenarios but the *UK10K* scenario, where we simulated 273 independent sets of 72 haplotypes) given an allele frequency trajectory *H*_*i*_ and we computed the *L* value for each pair of haplotypes. We can use that distribution of *L* values for a given allele frequency *H*_*i*_ to find the probability *P* (*L* ∈ *w*_*i*_ |*H*_*i*_) that *L* falls in a certain window *w*_*i*_. It is important to appreciate that these Monte Carlo simulations can include additional information about the recombination rate present in a particular region. Using the appropriate recombination rate is important because it changes the values of *L*.

The likelihood ℒ(4*Ns, f, D*|*L*) is found by integrating over the space of allele frequency trajectories that end at a frequency *f* in the present and have a selection coefficient 4*Ns*. One possible way to perform that integration step is to perform many simulations under the assumptions of the Poisson Random Field framework (Sawyer & Hartl 1992; Hartl *et al*. 1994) (*PRF*) and utilize rejection sampling to only keep those trajectories that end at a frequency *f* in the present. Under the *PRF* model, the number of mutations that enter the population each generation *i* have a Poisson distribution with mean *2NiμK* = Θ/2, where *N*_*i*_ is the population size in generation *i, μ* is the mutation rate per base and *K* is the number of sites being simulated. The sites are independent and the frequency of each mutation changes each generation following a Wright-Fisher model with selection. We could generate many allele frequency trajectories under this framework given a particular value of 4*Ns* and just keep those trajectories that end at a frequency of *f*. However, this is inefficient and computationally demanding, since the vast majority of allele frequency trajectories will not end at a frequency *f* in the present. And it is particularly more challenging if we wish to calculate ℒ(4*Ns, f, D*|*L*) for a grid of values of 4*Ns*. In the next section we show an alternative importance sampling approach we developed to perform an efficient integration over the space of allele frequency trajectories given 4*Ns* and *f*.

### Integration over the space of allele frequency trajectories using importance sampling

We used importance sampling to integrate over the space of allele frequency trajectories and calculate the likelihood ℒ(4*Ns, f, D*|*L*) over many different values of 4*Ns*. The efficient integration over the space of allele frequency trajectories is done using the importance sampling approach developed by Slatkin (2001) with a modification regarding the importance sampling distribution we use. Here, the “target” distribution *f* (*x*) = *P*(*H*_*i*_|*s, f*) are samples of allele frequency trajectories that end at a frequency *f* and have a selection coefficient *s*.

Following Slatkin (2001), we can define the trajectory *H*_*i*_ of a derived allele *a* as the number of copies of the allele *a* present each generation since the allele appeared in the population. Therefore, *H*_*i*_ = {*i*_*T*_, *i*_*T*−1_, *i* _*T*−2_, …, *i*_2_, *i*_1_ *i*_0_}, where *i*_*T*_ = 0 and *i*_*T*−1_ = 1. The effective population sizes at those times are *N* = {*N*_*T*_, *N*_*T*−1_, *N* _*T*−2_, …, *N*_2_, *N*_1_, *N*_0_}. The allele appears in generation *T-1*, where it has 1 copy in the population.

We define the fitness of the genotypes *AA, Aa* and *aa* as 1, 1+*s* and 1+2s, respectively. Under a Wright-Fisher model with selection, the probability of moving from *i*_*t*_ to *i*_*t*−1_ copies of the allele going forward in time is equal to:

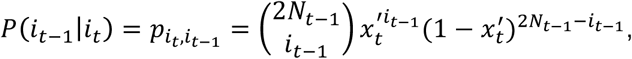

where

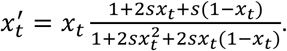

The frequency of the allele at generation *t* is 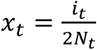.

As a “importance sampling” distribution *g(x)*, we use a very similar process to a Wright-Fisher neutral model. We start with the count *y* of the number of derived alleles *a* in the present based on a sample of *n* alleles. Estimating the frequency in generation 0 based on that sample of alleles is equal to the problem of estimating a probability based on binomial data. Therefore, we can follow Gelman *et al*. (2013) to state that the posterior density of the distribution of allele frequency 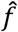 in generation 0 is distributed as: 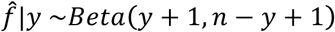. Based on the distribution of 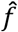, we can obtain the distribution of the number of alleles in generation 0, *i*_0_, just by multiplying 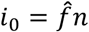 and rounding *i*_0_ to a discrete value. Then we can define the probability of having *i*_0_ alleles in generation 0 given that we sampled *y* derived alleles in a sample of *n* alleles as:

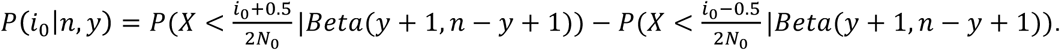

On the other hand, the probability that we obtain *y* derived alleles in a sample of *n* alleles given that the number of derived alleles in the population is *i*_0_ is:

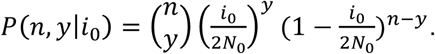

After we sample from that distribution, we move backwards in time assuming that the allele is neutral. Under this proposal distribution, if *i*_*t*−1_ = 1, then *i*_*t*_ can take any value from 0 to *2N*_*t*_. If *i*_*t*−1_ = 0 or *2N*_*t*_ then we stop the allele frequency trajectory. If *i*_*t*−1_ is bigger than 1 and smaller than *2N*_*t*_, then *i*_*t*_ can take any value from 1 to *2N*_*t*_. These three rules are used together to make sure that each trajectory going forward in time always goes from 0 to 1 copy of the allele.

Under the importance sampling distribution we use, the transition probabilities of going from *i*_*t*−1_ alleles in generation *t*-1 to *i*_*t*_ alleles in generation *i*_*t*_ is:

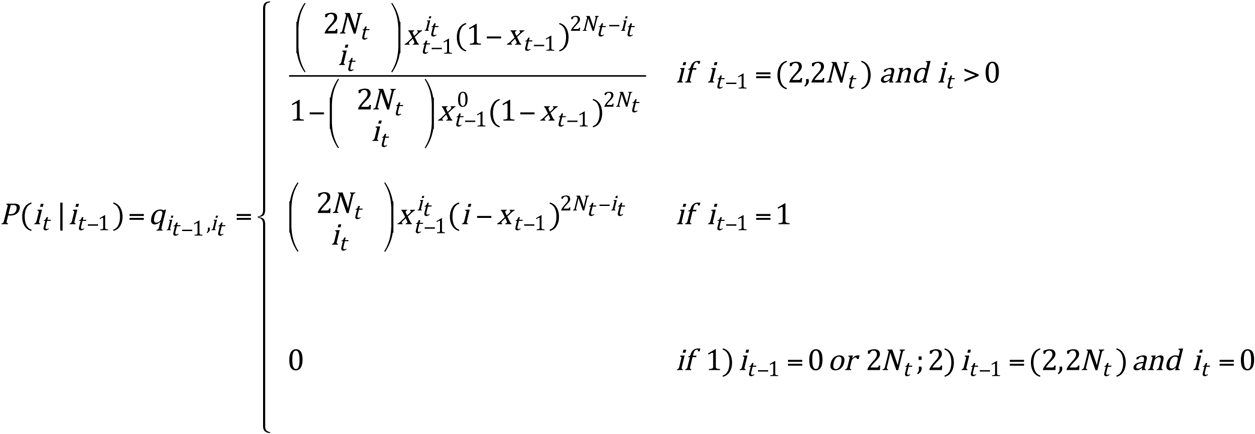

Where 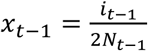. By generating an allele frequency trajectory with this importance sampling distribution, we can calculate the probability of any sample from this importance sampling distribution *g(x)*:

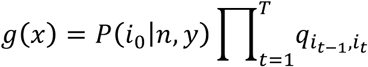

Finally, the probability of the whole allele frequency trajectory *H*_*i*_ going forward in time is then equal to:

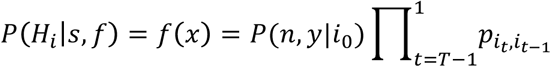

Now that we have defined how to sample allele frequency trajectories using our proposal distribution, we can compute the weight for every simulated allele frequency trajectory *H*_*i*_ from *g(x)* as 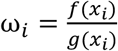. For some of the proposed trajectories under *g(x)*, the trajectory will end up at a frequency of 1 going backwards into the past, instead of 0. The value of ω_*i*_ for those trajectories is defined to be equal to 0.

The expected value that we wish to obtain with this problem is ℒ(4*Ns, f, D*|*L* ∈ *w*_*i*_). After generating *M* replicates using *g(x)*, we can compute that expected value under the importance sampling framework:

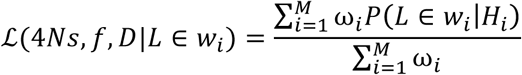

Using this approach, we can estimate *L*(4*Ns, f, D*|*L* ∈ *w* _*i*_) for different values of *s* using the same set of allele frequency trajectories generated from our importance sampling distribution. This alleviates the need to simulate a different set of allele frequency trajectories for each value of the selection coefficient *s* that we want to evaluate and follows the idea of a driving value (Fearnhead & Donnelly 2001). The proposal distribution *g(x)* is not necessarily optimal for every *s* value, but it is possible to verify if the distribution is reasonable based on the effective sample size (*ESS*) values (see Equation S1; Supplementary Methods). The *ESS* indicates the sample size used in a Monte-Carlo evaluation of the target distribution *f* (*x*) that is equivalent to the importance sampling approach estimate. Plots of the *ESS* values for the two main demographic scenarios explored are shown in the Supplementary Figures S18-S19. In every demographic scenario explored, we simulated 100,000 allele frequency trajectories to evaluate 401 values of 4*Ns* in discrete intervals from −200 to 200. The only values that we need to change to evaluate *L*(4*Ns, f, D*|*L* ∈ *w*_*i*_) are the importance sampling weights ω _*i*_, where we will change the value of *P* (*H* _*i*_| *s, f*)= *f* (*x*) depending on the value of the selection coefficient *s* evaluated.

Finally, given a set of values 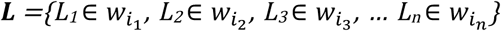, where *i*_*j*_ can take any value from 1 to *S*, we can estimate the composite likelihood of having that set of ***L*** values as:

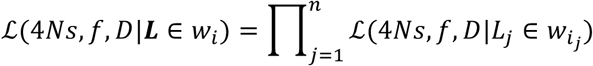

### Forward-in-time simulations to obtain mean allele frequency trajectories

We used *PReFerSim* (Ortega-Del Vecchyo *et al*. 2016) to obtain 10,000 allele frequency trajectories of a 1% frequency allele under the constant-size demography scenario for 5 different values of selection (4*Ns* = 0, −50, −100, 50, 100). To do those simulations, we performed many replicate simulations where the number of new mutations per generation follows a Poisson distribution with a mean equal to Θ/2 = 1,000. Those simulations were repeated until we obtained 10,000 alleles frequency trajectories where the present-day frequency *f* is equal to 1% in a sample of 4,000 chromosomes. We did the same procedure to obtain 10,000 allele frequency trajectories of a 1% frequency allele for 5 different values of selection (4*Ns* = 0, −50, - 100, 50, 100) in a population expansion and an ancient bottleneck scenario. The value of Θ/2 for the most ancestral epoch was set to 1,000 in the population expansion and the ancient bottleneck scenario.

In the case of the *UK10K* demographic scenario, we obtained 10,000 allele frequency trajectories of a 1% frequency allele for 5 values of selection (4*Ns* = 0, −25, −50, 25, 50). We performed many simulations using a Θ/2 value equal to 1,000 for the most ancestral epoch until we obtained 10,000 allele frequency trajectories. We sampled 7,242 chromosomes and retained those trajectories where *f* = 1% ± 0.05%.

### Connecting the distribution of fitness effects of variants at a particular frequency (*DFE*_*f*_) with the distribution of fitness effects of new mutations (*DFE*)

The distribution of fitness effects of variants at a particular frequency *DFE*_*f*_ in the population is related to the distribution of fitness effects of new mutations *DFE* defined by a set of *k* parameters ***ψ*** = {***ψ***_**1**_, ***ψ***_**2**_, ***ψ***_**3**_, …, ***ψ***_***k***_} by the following equation:

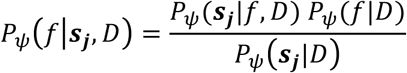

Where we can re-arrange the above equation to obtain:

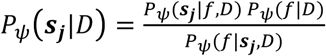

The events defined in that formula are:

*f*.- The allele has an x% sample allele frequency.

***s***_***j***_.- Allele has a selection coefficient 4*Ns* that falls in the interval [4*Ns*_*j-1*_, 4*Ns*_*j*_), where *s*_*j-1*_ and *s*_*j*_ define two different selection coefficients. *N* is the effective population size in the most ancestral epoch in the demographic scenario *D*.

*ψ*.- A set of *κ* parameters ***ψ*** = {*ψ*_1_, *ψ*_2_, *ψ*_3_, …, *ψ*_k_} that define the *DFE*.

*D*.- Demographic scenario.

*P*_*ψ*_(***s***_***j***_|*D*) defines the distribution of fitness effects of new mutations over a set of discrete bins when using the information contained across all non-overlapping intervals ***σ*** = *{[*4*Ns*_*0*_, 4*Ns*_*1*_*), [*4*Ns*_*1*_, 4*Ns*_*2*_*), [*4*Ns*_*2*_, 4*Ns*_*3*_*)…*, [4*Ns*_*b-1*_, 4*Ns*_*b*_*)} = {* ***s***_**1**_, ***s***_**2**_, ***s***_**3**_, …, ***s***_***b***_ *}* covering all 4*Ns* values from 0 to infinite. We defined the endpoints of the first *b*-1 intervals to be equal to 5(*i-*1) and 5*i*, where *i* takes values from 1 to *b* - 1, in all the analysis we performed with the exception of Supplementary Table S4. The last interval was set to be equal to [5*b*, ∞). Since *P*_*ψ*_ (***s***_***j***_|*D*) is independent of the demographic scenario *D*, then *P*_*ψ*_ (***s***_***j***_|*D*) = *P*_*ψ*_ (***s***_***j***_) because *D* does not impact the proportion of new variants in a selection interval *s*_*j*_. If we look at the information of all non-overlapping intervals ***σ***, *P*_*ψ*_ (***s***_***j***_|*f, D*) defines the distribution of fitness effects of variants at a particular frequency *DFE*_*f*_ over a set of discrete bins. As seen in the section *Testing inference of the distribution of fitness effects for variants found at a particular frequency (“DFE*_*f*_*”)*, we can use the ***L*** values to infer *DFE*_*f*_.

*P*_*ψ*_ (*f*|*D*) can be computed both in data and in simulations by measuring the proportion of variants at a certain frequency. Calculating *P*_*ψ*_ (*f*|*D*) in genomic data requires us to calculate the proportion of variants at a frequency *f*. That proportion must take into account all variants that have emerged during the demographic history *D*, including variants that have become fixed or have been lost. To calculate *P*_*ψ*_ (*f*|***s***_***j***_, *D*), we can make the assumption that all the mutations in the interval ***s***_***j***_ have very similar selection coefficients, which is more likely to be true when the interval is not very big. This probability can be found via forward-in-time simulations, where we simulate variants that have a selection coefficient contained in a certain interval ***s***_***j***_ in a particular demographic scenario *D*. Then, the proportion of variants in that simulation that have a *f* frequency in the present is equal to *P*_*ψ*_ (*f*|***s***_***j***_, *D*).

We calculate *P*_*ψ*_ (***s***_***j***_) for the first *b*-1 intervals using Equation 4. Then, for the last interval ***s***_***b***_ we use 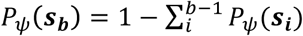. If 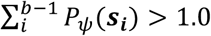 we set the probabilities 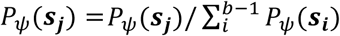 for the first *b*-1 intervals and *P*_*ψ*_ (***s***_***j***_) = 0 for the last interval *b*.

We tested Equation 4 on two different distributions of fitness effects (Figure 7 and Supplementary Figure S3). To perform those two tests we did simulations under the Poisson Random Field model using *PReFerSim* (Ortega-Del Vecchyo *et al*. 2016) to estimate *P*_*ψ*_ (*f*|***s***_***j***_, *D*). We did those simulations using the mouse distribution of fitness effects (Halligan *et al*. 2013) and the population expansion demographic model. Those calculations were done across 5,000 simulation replicates where the value of Θ/2 in the first epoch was set equal to 1,000. We sampled 4,000 chromosomes for each segregating site to calculate *f*.

When we estimated the distribution of fitness effects of new variants in the *UK10K* data, we estimated *P*_*ψ*_ (*f*|***s***_***j***_, *D*) by performing 1,000 replicate simulations under the inferred *UK10K* demographic model and the human distribution of fitness effects (Boyko *et al*. 2008). The value of Θ/2 in the first epoch of each simulation was set equal to 1,000. To mimic the properties of the *UK10K* data, we sampled 7,242 chromosomes for each segregating site. We calculated *P*_*ψ*_ (*f*|***s***_***j***_, *D*) by counting the proportion of variants in our 1,000 simulations that have a frequency *f* equal to *1%* ± 0.05%.

### Estimating *L* taking into account differences in local recombination rates in the *UK10K* dataset

Apart from being dependent on the strength of selection acting on the variants, the distribution of *L* surrounding each variant on the genome in the *UK10K* data is dependent on the local recombination rate *ρ*. We took into account the local recombination rate when inferring the distribution of fitness effects using the 273 nonCpG nonsynonymous 1% frequency variants. To do this, we used our importance sampling method to obtain the distribution of *L* given the selection coefficient, the inferred demographic scenario, and 21 different recombination rates. To select the 21 recombination rates, we used the results from a previously inferred recombination map (Kong *et al*. 2010). We took the 21 different percentile values (0^th^, 5^th^, …, 95^th^, 100^th^) from the distribution of 546 average recombination rates per base taken from the upstream and downstream 250 kb regions next to the 273 nonsynonymous 1% frequency variants. In the end, we generated 21 distributions for each selection value explored, each with a different recombination rate *ρ*_*j*_. Those 21 distributions of ℒ(4*Ns, f, D, ρ*_*j*_|*L* ∈ *w*_*j*_) were used to infer selection using the upstream and downstream regions from the nonCpG nonsynonymous 1% frequency variants. They were also used to infer the point estimate of 4*Ns* in the nonCpG synonymous 1% frequency variants. The ℒ(4*Ns, f, D, ρ*_*j*_|*L* ∈ *w* _*j*_) distribution used for each of the 546 regions is the one where the local recombination *ρ* is closer to *ρ*_*j*_.

We evaluated the accuracy of our method to infer selection under the inferred scaled *UK10K* demographic scenario using simulations. We mimicked the amount of information present in the *UK10K* data in each simulation replicate. Each simulation replicate contains 273 independent loci with 72 haplotypes containing the derived allele. The recombination rates, both to the left and right side of the loci, were equal to the average per base recombination rates in the 250 *kb* windows next to each locus in the data. We calculated *L* going to the left and right side of the focal loci, obtaining 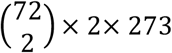 values for each simulation replicate (Supplementary Figure S8, S10).

## Data availability

The programs and data to reproduce every figure of the paper can be found in https://github.com/dortegadelv/HaplotypeDFEStandingVariation.

## Acknowledgements

We thank Christian Huber, Bernard Kim, Evan Koch, Charleston Chiang, Tanya Phung, Clare Marsden, Annabel Beichman, Jazlyn Mooney and Ying Zhen for useful discussions. We also thank Jair S. Garcia-Sotelo, Luis Alberto Aguilar Bautista, Alejandra Castillo and Carina Uribe for technical assistance. Infrastructure for computational analysis was in part provided by a CONACYT grant led by Alejandra Medina-Rivera [269449]. This work was supported by the UC MEXUS-CONACYT fellowship 213627 to D.O.-D.V, NIH grant R35GM119856 to K.E.L., and NIH grant R01HG007089 to J.N.

