## Supplementary Information for "Haplotype-based inference of the distribution of fitness effects"

### Supplementary Methods

#### Importance sampling

Broadly, the idea behind importance sampling approaches is that we have a "target" distribution $f(x)$ and we would like to take a large number of samples from that distribution. In this case, the "target" distribution $f(x)$ is the set of trajectories that end at frequency $f$ in the present. If we wanted to estimate expectations from that target distribution using a Monte Carlo method, we could take a set of samples $(x_1, x_2, \ldots, x_n)$ from the distribution $f(x)$ and then use the following equation:

$$E[f(x)] = \frac{\sum_{i=1}^{n} f(x_i)}{n}$$

In our case, sampling from this "target" distribution is complicated because when we simulate allele frequency trajectories going forward in time, the vast majority of them do not end at a frequency $f$ in the present. In the importance sampling framework, the idea is to choose a "Importance sampling distribution" $g(x)$ from which we can easily sample random values $X$. Then, making use of the equality $\frac{g(x)}{g(x)} = 1$, we can estimate the expected value of $f(x)$ by using the following equation:

$$E[f(x)] = \sum_{i=1}^{n} \frac{g(x_i)}{g(x_i)} f(x_i) = \sum_{i=1}^{n} \frac{f(x_i)}{g(x_i)} g(x_i)$$

Then, we define a variable $\omega_i = \frac{f(x_i)}{g(x_i)}$ and:

$$E[f(x)] = \sum_{i=1}^{n} \omega_i g(x_i)$$

In cases where either $f(x)$ or $g(x)$ are missing a normalizing constant so that their densities do not integrate to 1, we must employ a self-normalized importance sampling estimator (Robert & Casella 2010):

$$E[f(x)] = \frac{\sum_{i=1}^{n} \omega_i g(x_i)}{\sum_{i=1}^{n} \omega_i}$$

The selection of the “Importance sampling distribution” *g(x)* is critical in the importance sampling framework to accurately estimate the expected values of *f(x)*. Ideally, random variables simulated under *g(x)* could often be obtained by sampling under *f(x)*. A useful metric to assess the suitability of *g(x)* is the effective sample size *ESS*, which is equal to (Robert & Casella 2010):

$ESS = \frac{1}{\sum_{i=1}^{n} \underline{\omega_i}^2}$ (Equation *S1*)

Where:

$$\underline{\omega_i} = \frac{\omega_i}{\sum_{j=1}^{n} \omega_j}$$

The *ESS* indicate the sample size used in a Monte-Carlo evaluation of *f(x)* that is equivalent to the importance sampling approach estimate. The *ESS* takes values between 1 and *n*, where a higher value of the *ESS* indicates that more samples from *g(x)* are contributing to the estimate of the expected *f(x)*. This is a necessary, but not sufficient, condition to obtain an accurate estimate of the expected value of *f(x)* when using an importance sampling approach. Values of *ESS* close to 1 indicate that few replicates of *g(x)* are making a contribution of the expected value of *f(x)* and, therefore, the estimated expected value of *f(x)* is likely to not be accurate.

**Pairwise coalescent times $T_2$ given different values of selection**

We investigated properties of the distribution of $T_2$ given different strengths of selection. To do this, we discretized and compressed each of the allele frequency trajectories we obtained using *PReFerSim* according to a set of allele frequency boundaries (0.0, 0.001, 0.002, 0.003, 0.004, 0.005, 0.006, 0.007, 0.008, 0.009, 0.01, 0.0125, 0.015, 0.02, 0.03, 0.04, 0.05, 0.06, 0.07, 0.08, 0.09, 0.1, 0.15, 0.2, 0.3, 0.4, 0.5, 0.6, 0.7, 0.8, 0.9, 0.95, 0.96, 0.97, 0.98, 0.99, 0.995, 1.0) to reduce the computational time needed to simulate haplotypes under the structured coalescent model with *mssel*. We then used each compressed allele frequency trajectory to estimate the distribution of pairwise coalescent times $T_2$ between a pair of haplotypes containing the allele changing in frequency. For each compressed allele frequency trajectory, we estimated the probability of coalescing at a time $t$ as:

$$P(T_2 = t) = \left[\prod_{x=1}^{t-1}\left(1-\frac{1}{N_{a_x}}\right)\right]\frac{1}{N_{a_t}}$$

Where $N_{a_x}$ denotes the number of chromosomes that have the derived allele at generation $x$. Additionally, due to the way we compressed the allele frequency trajectories, where the allele frequency at the time that the allele emerges is not equal to $1/N_{a_x}$, the probability of $T_2$ in the generation $j$ where the allele appears is equal to $1-\sum_{t=1}^{j-1} P(T_2 = t)$. We averaged the probabilities of $T_2$ over all the simulated allele frequency trajectories given a particular value of $4Ns$ to obtain the distribution of $T_2$ given a value of $4Ns$.

**_ABC_ algorithm used to infer the demographic scenario consistent with haplotypic patterns at 1% frequency synonymous variants**

Our *ABC* algorithm has the following steps:

Data preparation:

1) Define the frequency $f$ of the variants that will be used to calculate the distribution of the haplotypic lengths $L$. We will use that distribution $L$ to infer the demographic history that better explains the haplotypic patterns observed in the variants at those frequencies. We will infer a demographic history for variants at a frequency $f = 1\% \pm 0.05\%$.
2) Out of the 3,781 individuals present in the phased *UK10K* haplotype reference panel, we selected the 3,621 individuals that had European ancestry, along with a set of individuals that were not related to other individuals in the panel, as previously defined in the original *UK10K* study (Walter *et al.* 2015).
3) We estimated the frequency of every variant present in the phased haplotype panel.
4) To identify the synonymous variants, we used the functional annotations for each variant from Ensembl 75 Variant Effect Predictor, as reported in the vcf file with allele frequencies available from the *UK10K* website (https://www.uk10k.org/data.html). Since some variants annotated as a synonymous mutation can possess more than one functional annotation, we only considered a variant to be synonymous if it did not possess another annotation that had a higher impact (HIGH or MODERATE) as defined in https://www.ensembl.org/info/genome/variation/prediction/predicted_data.html . . Variants with a high or moderate impact are those annotated to be transcription ablation, splice acceptor variants, splice donor variants, stop gained, frameshift, stop lost, start

lost, transcript amplification, inframe insertion, inframe deletion, missense, or protein altering variant.

5) We used the annotations from the 1000 Genomes Project phase 3 (Auton *et al.* 2015) to define the ancestral allele of each variant. This also allows us to define the derived allele of every site.
6) Depending on the frequency $f$ of the variants being investigated, retain all the $Y$ synonymous variants $V = \{V_1, V_2, \ldots, V_Y\}$ that: 1) Have a derived allele frequency in the interval $1\% \pm 0.05\%$ and 2) are more than 5 Mb away from the centromeres or telomeres. In the end we retained 152 synonymous variants.
7) For every variant $V_i$, take the $K_i$ haplotypes $h_{Ki} = \{h_1, h_2, \ldots, h_{Ki}\}$ containing the derived allele. Then, take the $h_{Ki}$ haplotypes and remove the sites where there is a singleton variant. After removing those sites, calculate the haplotypic length $L$ for every possible pair of haplotypes in $h_{Ki}$. The *UK10K* haplotype panel does not contain singleton variants present in the sample of 7,242 chromosomes. Therefore, the calculation of $L$ in this step is done ignoring singleton sites, either because they were not present in the original *UK10K* panel or because we remove them from the sample of $h_{Ki}$ haplotypes. Since there are $P_i = K_i*(K_i-1)/2$ possible pairs of haplotypes, after performing this step we will end up with $P_i$ values of $L$ for each variant $V_i$. The total number of $L$ values will be equal to $P = \sum_{i=1}^{S} P_i$.
8) We define 6 nonoverlapping windows $\boldsymbol{W} = \{w_1, w_2, w_3, w_4, w_5, w_6\} = \{ (0,50000], (50000, 100000], (100000,150000], (200000,250000], (250000, \infty)\}$. Then, we calculate $P(L \in w_i)$ using the distribution of $P$ values of $L$ obtained in the previous step. The distribution $D_{w_i} = P(L \in w_i)$ across the 6 $i$ windows represents the summary statistic we will use to infer demography in our ABC algorithm.

*ABC* algorithm:

We constructed demographic models for the synonymous variants at a $1\% \pm 0.05\%$ frequency in the population. A graphical representation of the model is presented in Supplementary Figure S5, along with the prior distributions for the demographic parameters of the models.

Our *ABC* approach follows this procedure:

1) Define the frequency $X = 1\% \pm 0.05\%$ of the alleles that will be investigated.
2) Draw a random value from each prior distribution of demographic parameters.

3) Using *PReFerSim*, simulate $Y$ random allele frequency trajectories $H_i$ where the allele ends at a frequency $X$=$1\% \pm 0.05\%$ in the present based on the parameter values sampled from the prior distribution. To reduce the running time, in these simulations we reduced all population sizes and times of population size changes by a factor of five.
4) Simulate $K$ haplotypes containing the derived allele for each simulated allele frequency trajectory obtained in the past step using *mssel*. We defined $K$ to be equal to $72$. $K$ is close to the product of $NX$, where $N$ is the total number of haplotypes in the *UK10K* dataset ($7{,}242$). After this step, $Y$ datasets with $K$ derived haplotypes were simulated.
5) For each to the $Y$ datasets, calculate the value of $D'_{w_i} = P(L \in w_i)$ by taking all possible pairs of haplotypes with the derived allele. We defined the values of $L$ using the same windows $\boldsymbol{W}$ employed in the *UK10K* dataset. The *UK10K* dataset does not contain singleton sites. To mimic this aspect from the data, the calculation of the $L$ values is done ignoring singleton sites. In the end, a distribution of $L$ values is obtained, where the total number of $L$ values is equal to $P = S\frac{K(K-1)}{2}$.
6) Calculate $\alpha = \sum_{i=1}^{6} |D'_{w_i} - D_{w_1}|$
7) Go back to 2) until we have sampled $50{,}000$ values for each demographic parameter from the prior distributions.
8) Retain the $100$ simulations where the value of $\alpha$ is closest to the one seen in the data. The values obtained for those parameters in those $100$ simulations define the posterior distributions of those parameters. The point estimates of each parameter were defined as the median of the posterior distribution.

The values of $\alpha$ in the $100$ retained simulations went from $0.0128$ to $0.0284$, indicating that we had a good match between the data and the $100$ retained simulations. Additionally, we performed $100$ simulations using the point estimates of the demographic parameters, and found that the average value of $\alpha$ was equal to $0.029$ using those $100$ simulations.

A recent study found that recent migration from Africa to Great Britain is an important phenomenon for alleles with a current frequency smaller than $0.35\%$ in the Great Britain population, as assessed using the *UK10K* dataset (Platt & Hey 2017). That migration altered the age distribution of alleles with smaller frequencies than $0.35\%$ in the *UK10K* dataset and needs to be modeled to fit the distribution of ages in those rare alleles. Therefore, it is possible that we would need to fit a scenario with more than one deme when looking at variants with a frequency smaller than $0.35\%$. That same study (Platt & Hey 2017) found that the *UK10K* > 0.35% frequency variants have primarily not been introduced in the Great Britain population due to recent migration. Those > 0.35% frequency alleles appeared before the divergence of African and European populations, implying also that alleles at higher frequencies than 0.35% have not been recently introduced to Great Britain from Africa.

#### Bootstrap confidence intervals

We used a bootstrap approach to estimate the 95% confidence intervals of our estimate of the selection coefficient $s$. We resampled each of the 273 variants with replacement, with their respective $L$ values, and we estimated the value of selection using this distribution of $L$ values. This process was repeated using a sample of 100 bootstrap replicates.

We used the same bootstrap approach to estimate the shape and scale parameters of a compound distribution of fitness effects. The variation across 100 bootstrap replicates is shown in Supplementary Figure S14.

#### Estimation of $P_{\psi}(f|D)$ in the *UK10K* dataset

To estimate $P_{\psi}(f|D)$, we used a ratio of two values:

1) Numerator: The number of nonsynoymous nonCpG variants that are more than 5 Mb away from centromeres or telomeres. This is 273.
2) Denominator: We calculated the total number of mutations in the *UK10K* demographic scenario (see Supplementary Figure S5), including the alleles that became fixed or extinct from the population. For each generation in our demographic scenario, the total number of mutations is equal to: $2 \times N \times l \times \mu$. Where $N$ is the effective population size in that generation. $l$ is the total number of nonCpG nonsynonymous possible variants that are more than 5 Mb away from centromeres or telomeres (26,368,474). We used the Ensembl 75 Variant Effect Predictor 93.3 to aid in our calculation of $l$. A variant was annotated as nonsynonymous if it possessed a nonsynonymous annotation in any of the transcripts inspected and it did not possess a variant that gave a “high” impact in any transcript as explained in https://www.ensembl.org/info/genome/variation/prediction/predicted_data.html. Variants with a high impact are those annotated to be transcription ablation, splice acceptor variants, splice donor variants, stop gained, frameshift, stop lost, start lost or transcript

amplification. Since each nonCpG site on the genome can mutate to three alternative nucleotides, each nonCpG site can contribute a number of $0$ to $3$ possible alternative nonsynonymous variants. $\mu$ is the mutation rate, which we set to $1.5 \times 10^{-8}$ (Ségurel *et al.* 2014). We sum the total number of mutations across all the generations to obtain the total number of mutations in the *UK10K* demographic scenario.

We show our estimates of $P_{\psi}(\boldsymbol{s_j})$ over 3 different $\boldsymbol{s_j}$ intervals in Figure 9 using the estimate of $P_{\psi}(f|D)$ described above. We compare our estimate of $P_{\psi}(\boldsymbol{s_j})$ with previous estimates (Boyko *et al.* 2008; Kim *et al.* 2017).

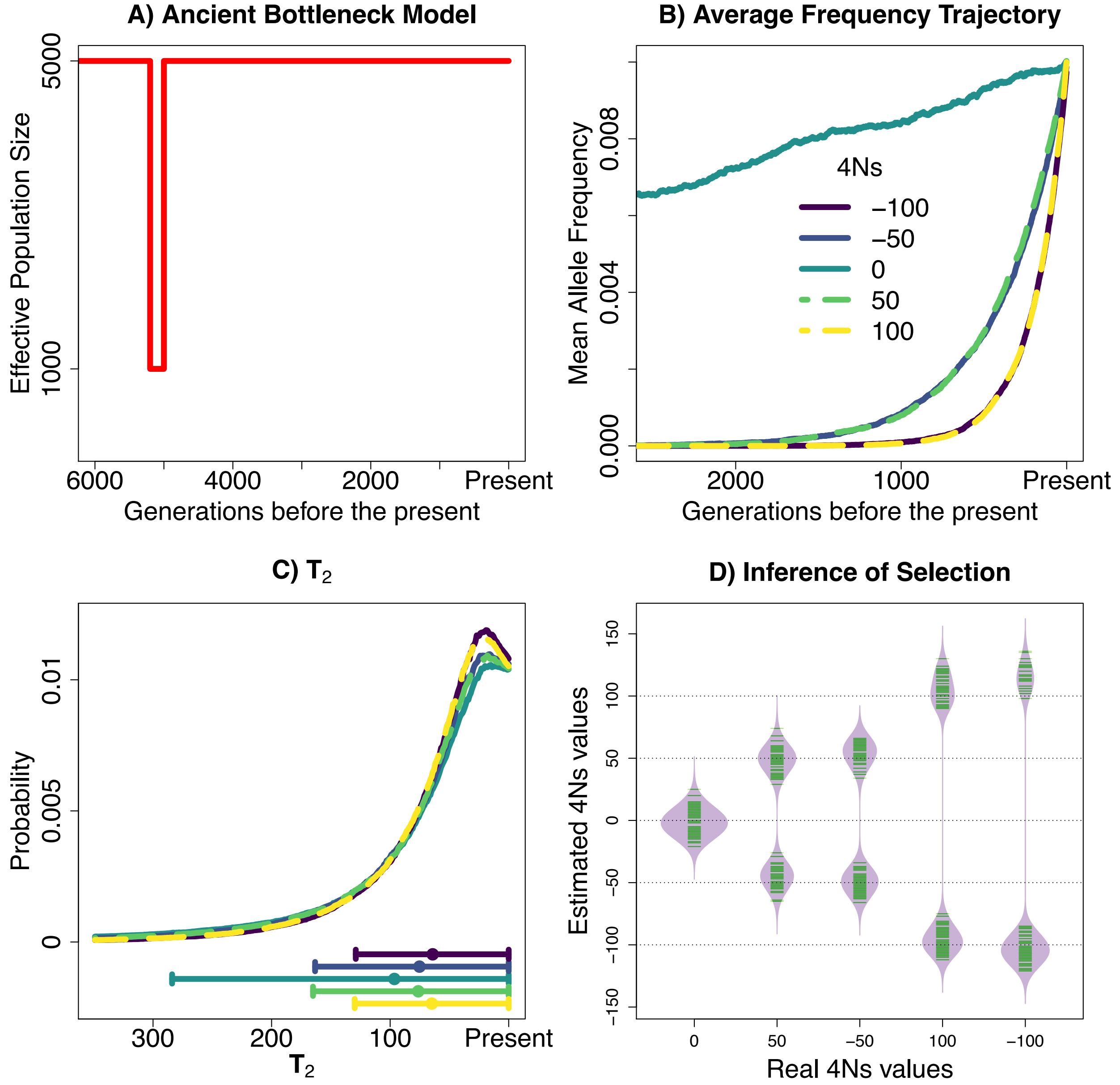


**Supplementary Figure S1.- Properties of alleles sampled at a 1% frequency under different strengths of natural selection in a scenario with a bottleneck that took place 5,000 generations ago.**

A) Demographic model analyzed. B) Mean allele frequency at different times in the past, in units of generations. C) Probability distribution of pairwise coalescent times $T_2$. The dot and whiskers represent the mean value of the distribution and the two whiskers extend at both sides of the mean until max(mean +- s.d. ,0).and D) Estimation of the strength of selection using 100 simulations for each $4Ns$ value, each with $10{,}000$ $L$ values. The green lines in D) indicate estimated values of $4Ns$. The recombination rate in the simulated 250 kb region for the most recent epoch was set equal to $\rho = 4Nr = 100$ and the mutation rate was set equal to $\theta = 4Nu =$ 100.

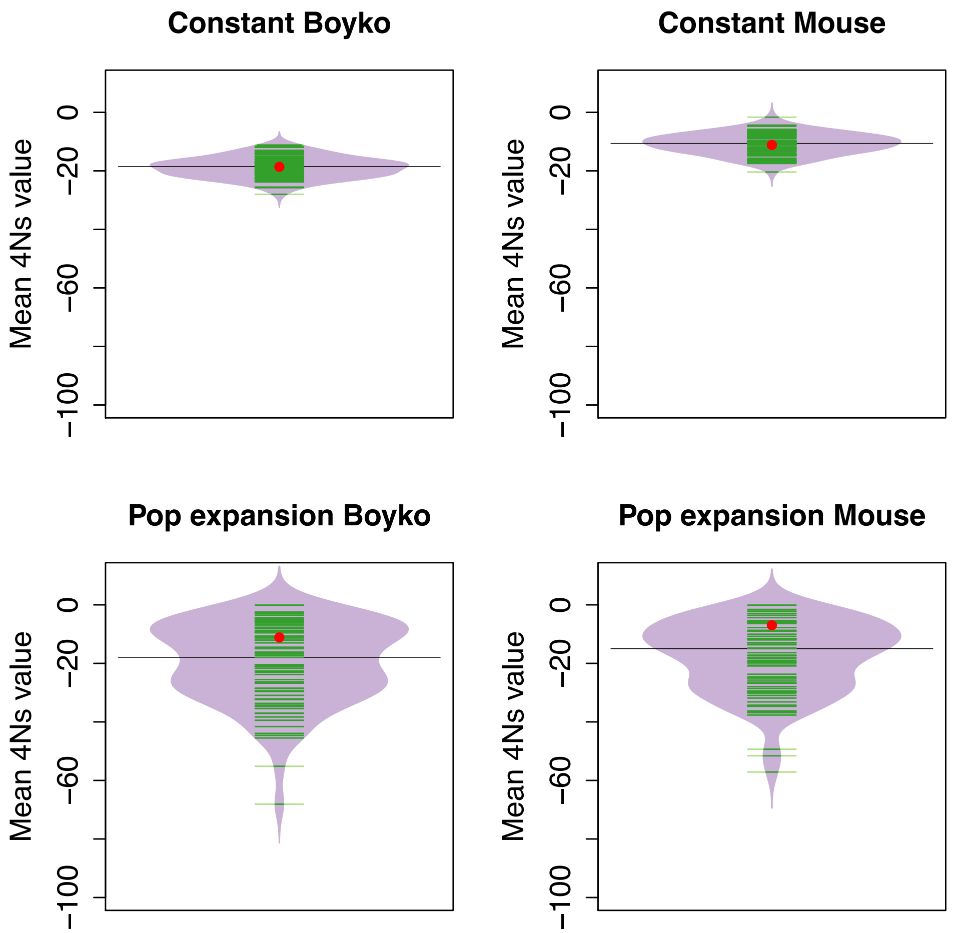


**Supplementary Figure S2.- Estimation of the mean $4Ns$ values for variants segregating at a $1\%$ frequency in simulated data.**
The beanplots show the distribution of the estimated mean $4Ns$ values based on the $DFE_f$ estimated on $100$ simulation replicates done with $100{,}000$ $L$ values. The red dots show the actual mean $4Ns$ value from $50{,}000$ $1\%$ frequency variants from each particular $DFE$ and demographic model employed. The green lines indicate estimated values of $4Ns$ across replicates. The median value of the estimates of $4Ns$ is shown with a solid line.

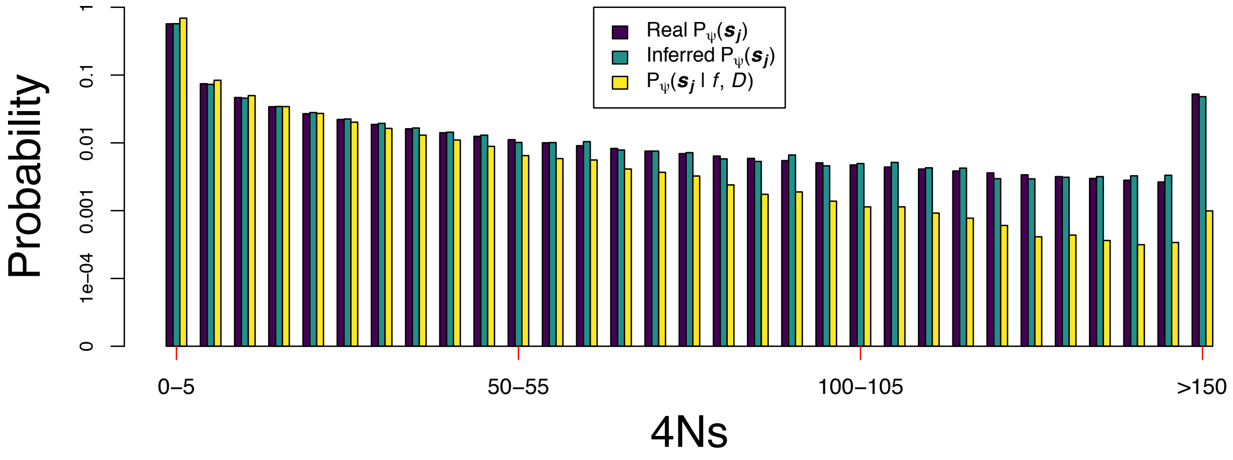


**Supplementary Figure S3.- Inference of the distribution of fitness effects of new mutations from the distribution of fitness effects of variants at a certain frequency in deleterious variants.**

The $DFE$ follows a gamma distribution with shape and scale parameters equal to $0.184$ and $79.966$, respectively. The results are for a population expansion demographic scenario where the population grows from $5{,}000$ to $50{,}000$ individuals in the last $100$ generations (see also Figure 4A). 'Real $P_{\psi}(\boldsymbol{s_j})$' refers to the probability of having a $4Ns$ value in a certain interval $\boldsymbol{s_j}$ given the distribution of fitness effects of new mutations with parameters $\psi$. $P_{\psi}(\boldsymbol{s_j}|f, D)$ is the probability of having a $4Ns$ value in an interval $\boldsymbol{s_j}$ given the distribution of fitness effects $DFE$ with parameters $\psi$ and the demographic scenario $D$ in $1\%$ frequency variants. We calculated $P_{\psi}(\boldsymbol{s_j}|f, D)$ from a set of $\sim 40{,}000$ $4Ns$ $1\%$ variants obtained via *PReFerSim* simulations under the $DFE$ and the population expansion scenario. 'Inferred $P_{\psi}(\boldsymbol{s_j})$' is an estimate of the probability of having a $4Ns$ value in a certain interval $\boldsymbol{s_j}$ given the distribution of fitness effects of new mutations. That probability was estimated using $P_{\psi}(\boldsymbol{s_j}|f, D)$ and Equation $4$. The selection coefficient $s$ refers exclusively to the action of deleterious variants in this plot.

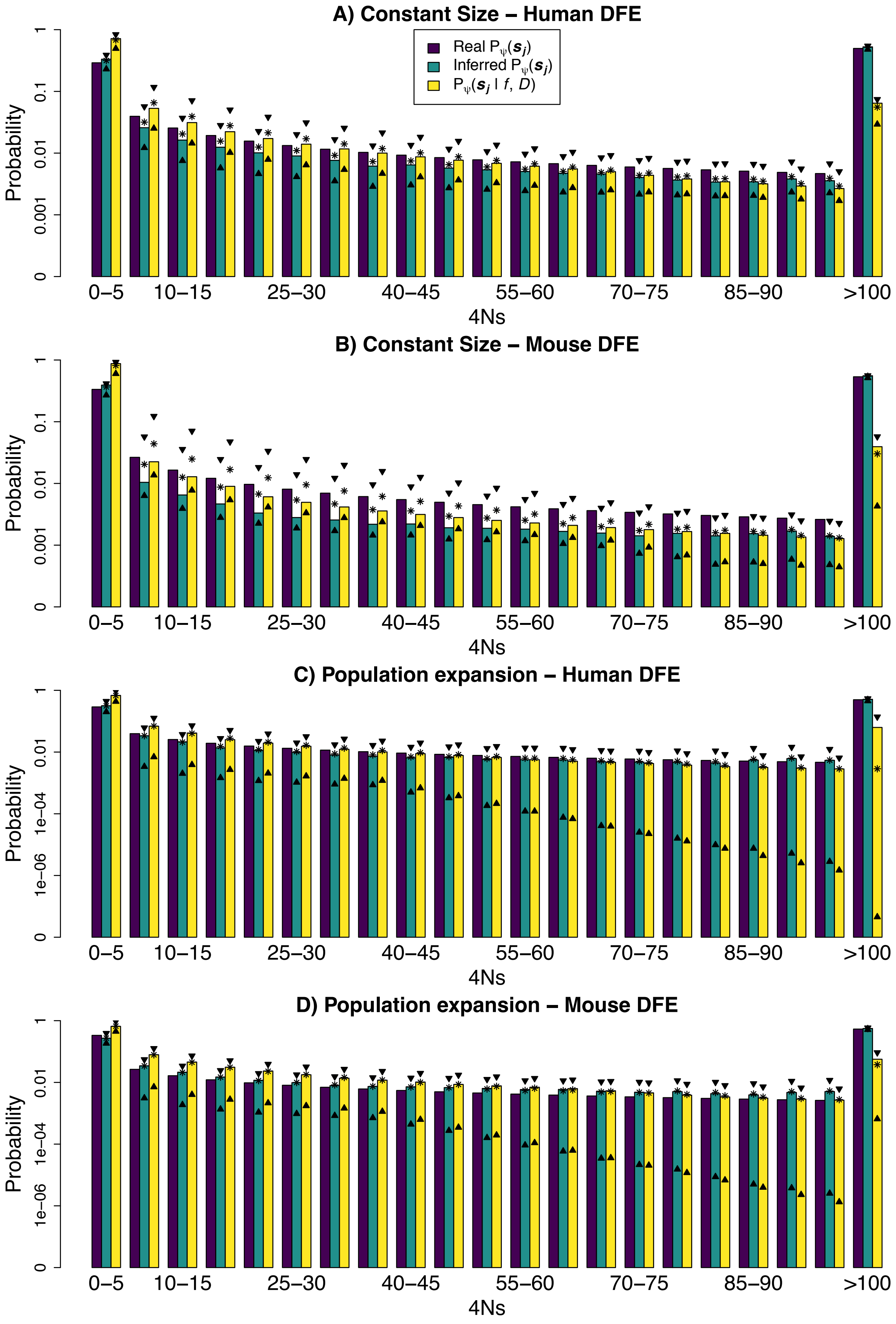
A) Constant Size – Human DFE
Real Pψ(sj)
Inferred Pψ(sj)
Pψ(sj | f, D)
Probability
0
0.001
0.01
0.1
1
0–5
10–15
25–30
40–45
55–60
70–75
85–90
>100
4Ns
B) Constant Size – Mouse DFE
C) Population expansion – Human DFE
1e−06
1e−04
D) Population expansion – Mouse DFE

236

**Supplementary Figure S4.- Inference of the distribution of fitness effects of new mutations from the distribution of fitness effects of variants in $1\%$ frequency variants.** 'Real $P_\psi(\boldsymbol{s_j})$' is the proportion of variants in a certain $\boldsymbol{s_j}$ interval based on the parameters $\psi$ that define the distribution of fitness effects of new variants $DFE$. The bars on '$P_\psi(\boldsymbol{s_j}|f, D)$' and 'Inferred $P_\psi(\boldsymbol{s_j})$' represent the median value of those respective estimated probabilities across $100$ simulation replicates employing $100{,}000$ $L$ values. The two triangles shown in each $\boldsymbol{s_j}$ interval denote the $5\%$ and $95\%$ percentile of the 'Inferred $P_\psi(\boldsymbol{s_j})$' or '$P_\psi(\boldsymbol{s_j}|f, D)$' probabilities estimated across $100$ simulation replicates. The mean values of '$P_\psi(\boldsymbol{s_j}|f, D)$' and 'Inferred $P_\psi(\boldsymbol{s_j})$' from 100 simulations are shown with an asterisk.

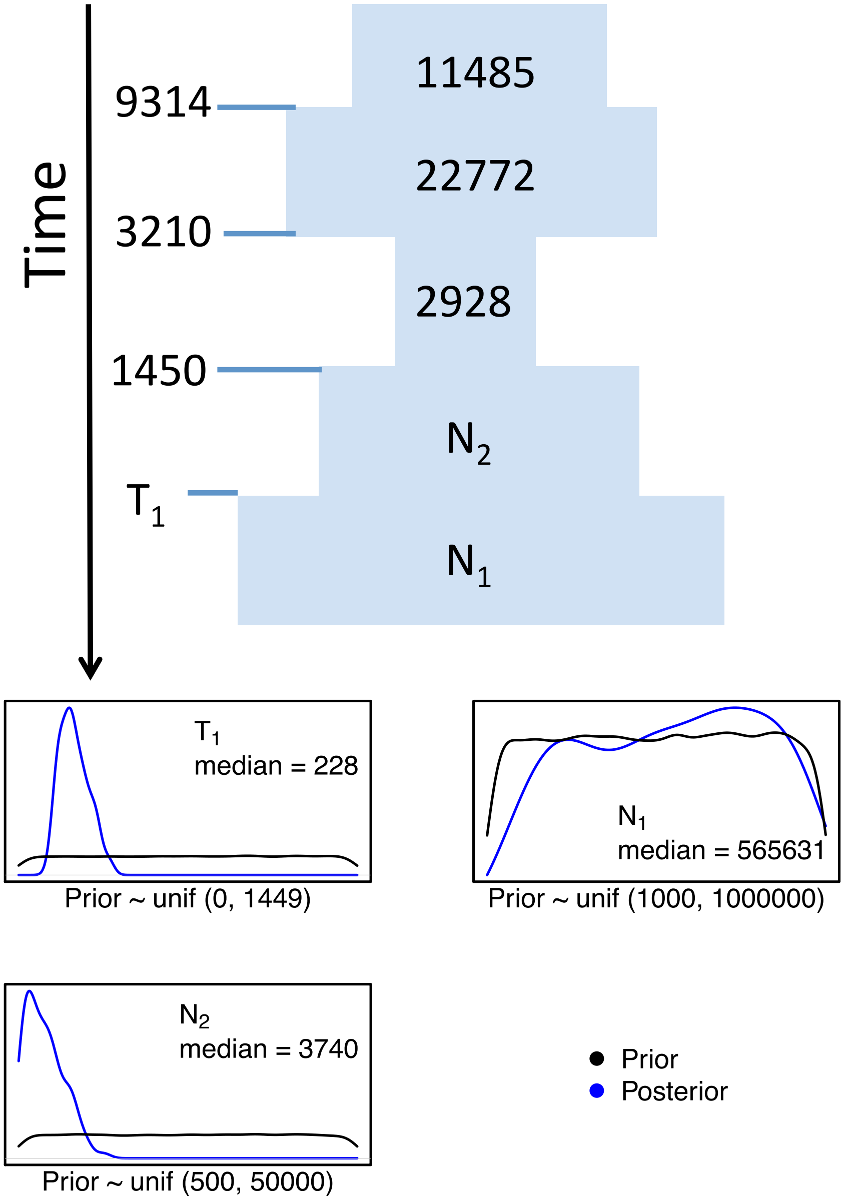


**Supplementary Figure S5.- Inference of demographic models that fit the $L$ distribution at 1% frequency synonymous variants in the *UK10K* dataset.** The plots below show the prior and posterior distribution for the demographic parameters of each of those models.

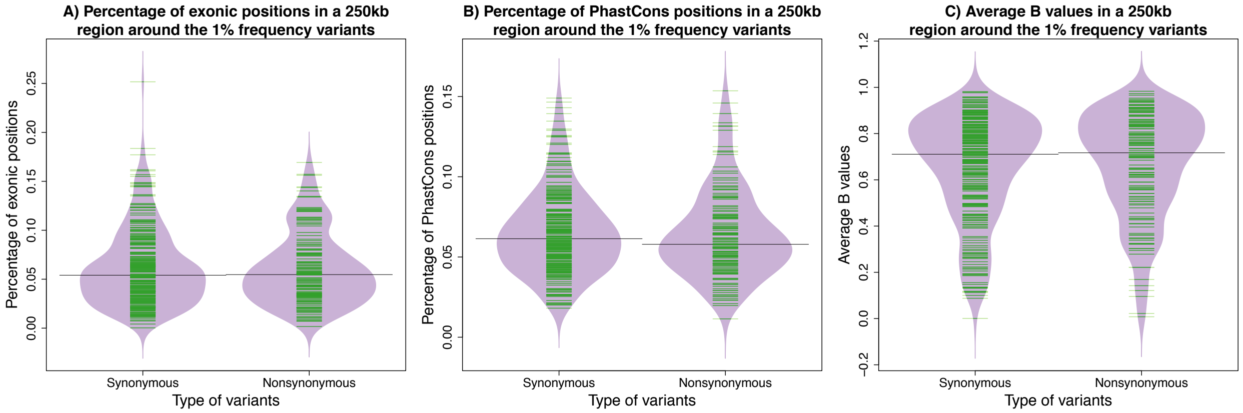


**Supplementary Figure S6.- Percentage of exonic positions, percentage of *PhastCons* element positions and average *B* values in the 250 kb upstream and downstream region of the inspected 1% frequency variants.**

A) The proportion of exonic positions for 1% frequency synonymous and nonsynonymous variants has a non-significant difference in mean values (Syn = 6.22%; Nonsyn = 6.20%; Mann-Whitney U test for mean differences p –value = 0.876), similar medians (Syn = 5.44%; Nonsyn = 5.49%) and a non-significant difference in their variances (Syn = 0.143%; Nonsyn = 0.148%; Brown-Forsythe test for the difference in variances p-value = 0.7895), respectively.

B) The proportion of positions in a *PhastCons* element for 1% frequency synonymous and nonsynonymous variants have a very similar mean (Syn = 6.58%; Nonsyn = 6.40%; Mann-Whitney U test for mean differences p –value = 0.299), median (Syn = 6.10%; Nonsyn = 6.00%) and a non-significant difference in their variances (Syn = 0.073%; Nonsyn = 0.072%; Brown-Forsythe test for the difference in variances p-value = 0.9284).

C) The average B values for 1% frequency synonymous and nonsynonymous variants display a very similar mean (Syn = 65.74%; Nonsyn = 65.07%; Mann-Whitney U test for mean differences p –value = 0.605), similar medians (Syn = 69.67%; Nonsyn = 70.81%), and a non-significant difference in their variances (Syn = 4.86%; Nonsyn = 4.99%; Brown-Forsythe test for the difference in variances p-value = 0.7298).

The exonic positions were taken from the UCSC table browser (Karolchik *et al.* 2003) using the following Table Browser controls: Clade = Mammal; Genome = Human; Assembly = hg19; Group = Genes and Gene Predictions; Track = UCSC Genes; Table = known gene; Region = Genome.

The positions of the PhastCons elements were obtained with these Table Browser controls: Clade = Mammal; Genome = Human; Assembly = hg19; Group = Comparative Genomics; Track = Conservation; Table = 100 Vert. El (phastConsElements100way); Region = Genome.

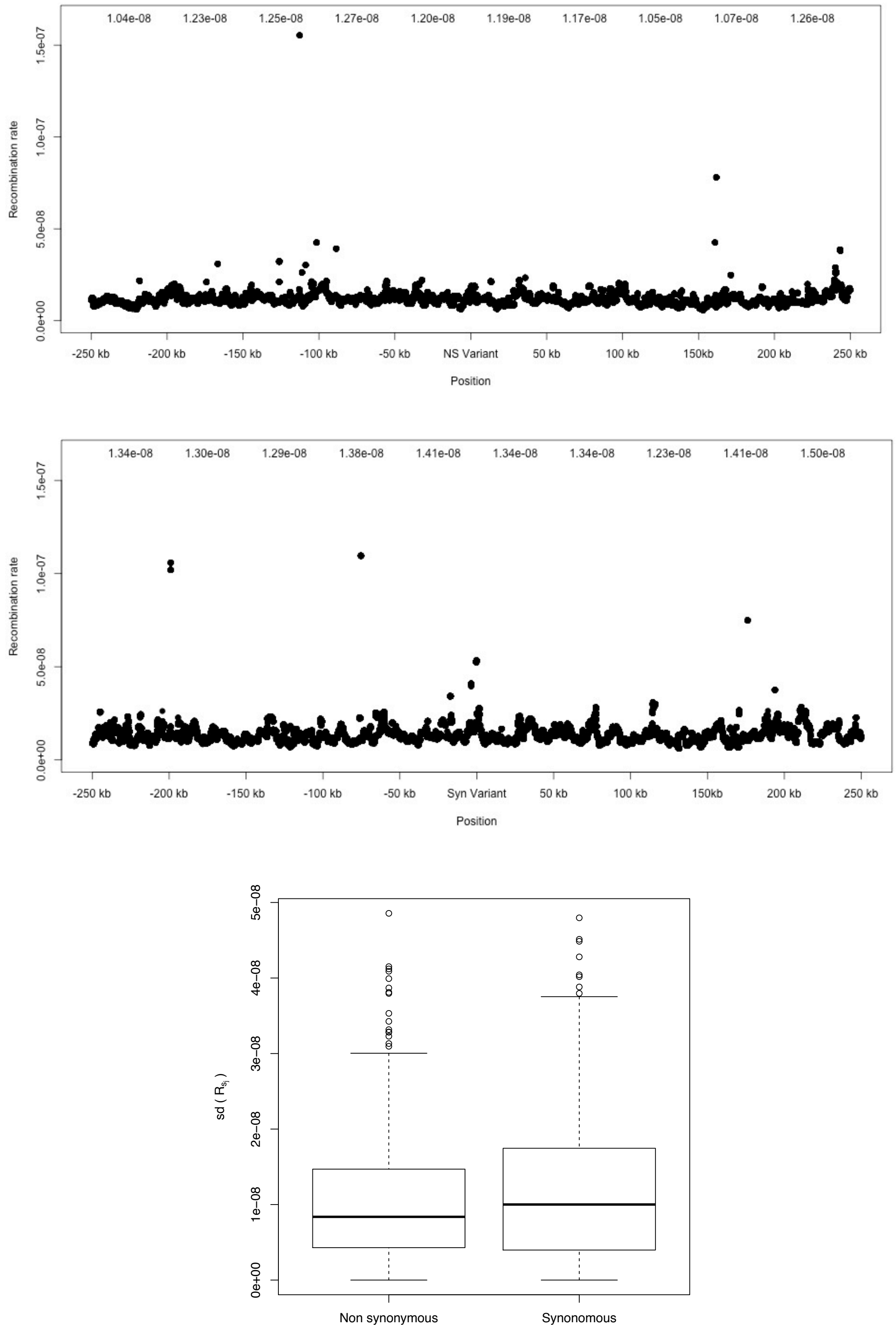


**Supplementary Figure S7.- Distribution of recombination rate values in the vicinity of the synonymous and nonsynonymous sites.**

Upper panel.- Mean recombination rate values 250 kb downstream and 250 kb upstream of the 273 1% nonsynonymous variants analyzed in the *UK10K* dataset. The black dots indicate the mean recombination values at each position upstream or downstream of the 273 nonsynonymous variants analyzed. The numbers shown at the top indicate the mean recombination values in each of the 10 $\boldsymbol{s_j}$ windows = {(-250kb, -200 kb), (-200kb, -150 kb), (-150kb, -100 kb), (-100kb, -50 kb), (-50kb, 0 kb), (0 kb, 50 kb), (50 kb, 100 kb), (100 kb, 150 kb), (150 kb, 200 kb), (200 kb, 250 kb)}.

Middle panel.- Mean recombination rate values 250 kb downstream and 250 kb upstream of the 152 1% synonymous variants analyzed in the *UK10K* dataset. The black dots indicate the mean recombination values at each position upstream or downstream of the 152 synonymous variants analyzed. The numbers shown at the top indicate the mean recombination values in each of the 10 $\boldsymbol{s_j}$ windows = {(-250kb, -200 kb), (-200kb, -150 kb), (-150kb, -100 kb), (-100kb, -50 kb), (-50kb, 0 kb), (0 kb, 50 kb), (50 kb, 100 kb), (100 kb, 150 kb), (150 kb, 200 kb), (200 kb, 250 kb)}.

Bottom panel.- In the left boxplot we plot the standard deviation in the 5 mean recombination rates $\boldsymbol{R_{sj}}$ of the 5 windows downstream {(-250kb, -200 kb), (-200kb, -150 kb), (-150kb, -100 kb), (-100kb, -50 kb), (-50kb, 0 kb)} for each one of the 273 1% nonsynonymous variants analyzed. We also plot the standard deviation in the 5 mean recombination rates $\boldsymbol{R_{sj}}$ of the 5 windows upstream (0 kb, 50 kb), (50 kb, 100 kb), (100 kb, 150 kb), (150 kb, 200 kb), (200 kb, 250 kb)}.

The right boxplot shows the same analysis for the 152 1% synonymous variants.

We conclude that in the 250 kb region upstream and downstream of the inspected synonymous and nonsynonymous variants there is variation in the recombination rates, however the variation is not very large since the standard deviation is similar to the mean recombination rate across the whole region. Further, there is no clear positional effect such that recombination rates increase or decrease markedly with distance from the focal SNP. Finally, we observe the recombination rates are similar (though significantly different) between non-synonymous and synonymous sites (mean recombination rate within 250kb for nonsynonymous = 1.18 $x$ $10^{-8}$; mean recombination rate within 250kb for S = 0.74 $x$ $10^{-8}$; p-value Mann-Whitney U Test < 2.2 $x$ $10^{-16}$).

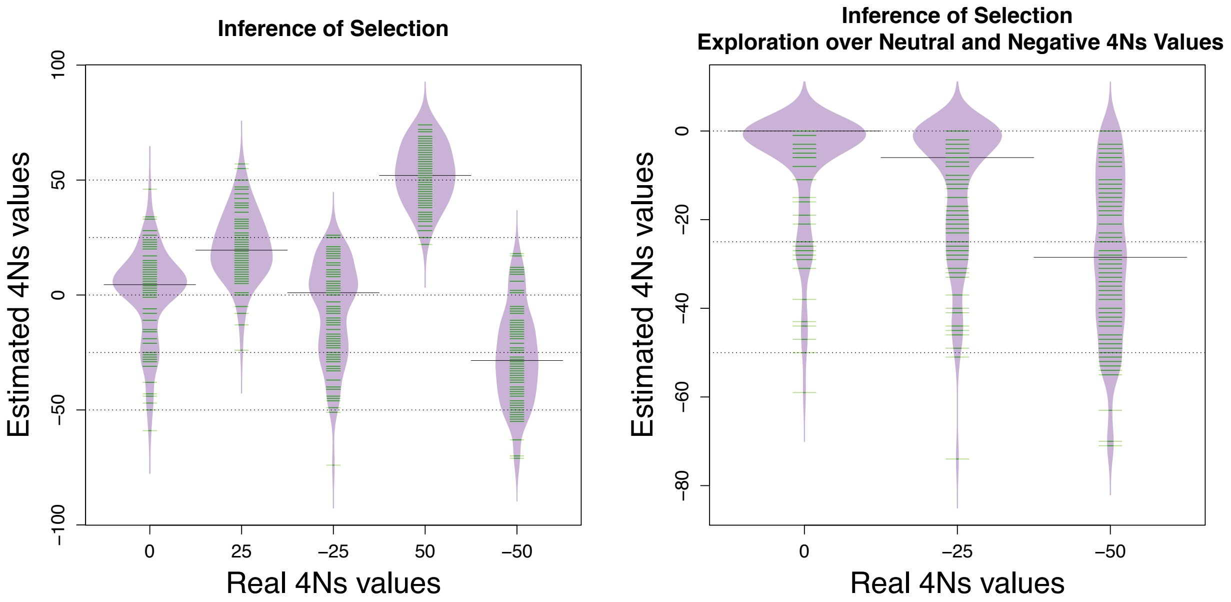


**Supplementary Figure S8.- Estimation of the strength of natural selection under the demographic model inferred from the scaled *UK10K* dataset on simulations.** Each simulation replicate contains $273$ independent loci with $72$ haplotypes containing the derived allele. We calculated $L$ going upstream and downstream of the focal loci, obtaining $\binom{72}{2} \times 2 \times 273$ $L$ values for each simulation replicate.

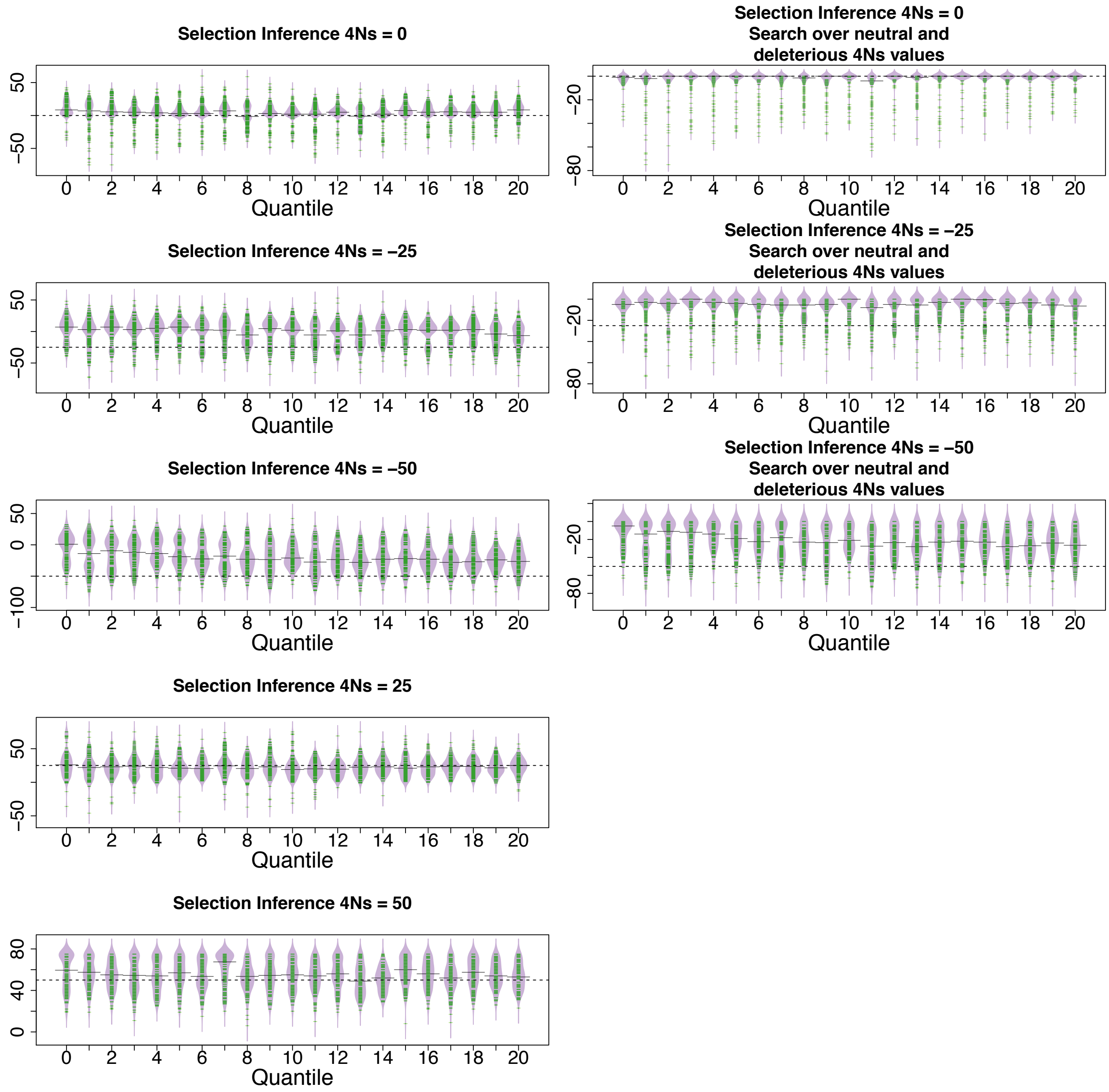


**Supplementary Figure S9.- Inference of selection under the scaled *UK10K* demographic scenario for 5 different values of selection and 21 different recombination rates.** We inferred selection across 100 simulation replicates for each of the 21 recombination rates explored. The 21 different recombination rates used are the 21 different percentile values ($0^{th}$, $5^{th}$, ..., $95^{th}$, $100^{th}$ ) from the distribution of 546 average recombination rates per base taken from the upstream and downstream 250 kb regions next to the 273 nonsynonymous 1% frequency variants. Each simulation replicate contains 273 independent loci with 72 haplotypes containing the derived allele and a fixed recombination rate for all the loci. We calculated $L$ going upstream and downstream of the focal loci, obtaining $\binom{72}{2} \times 2 \times 273$ $L$ values for each simulation replicate.

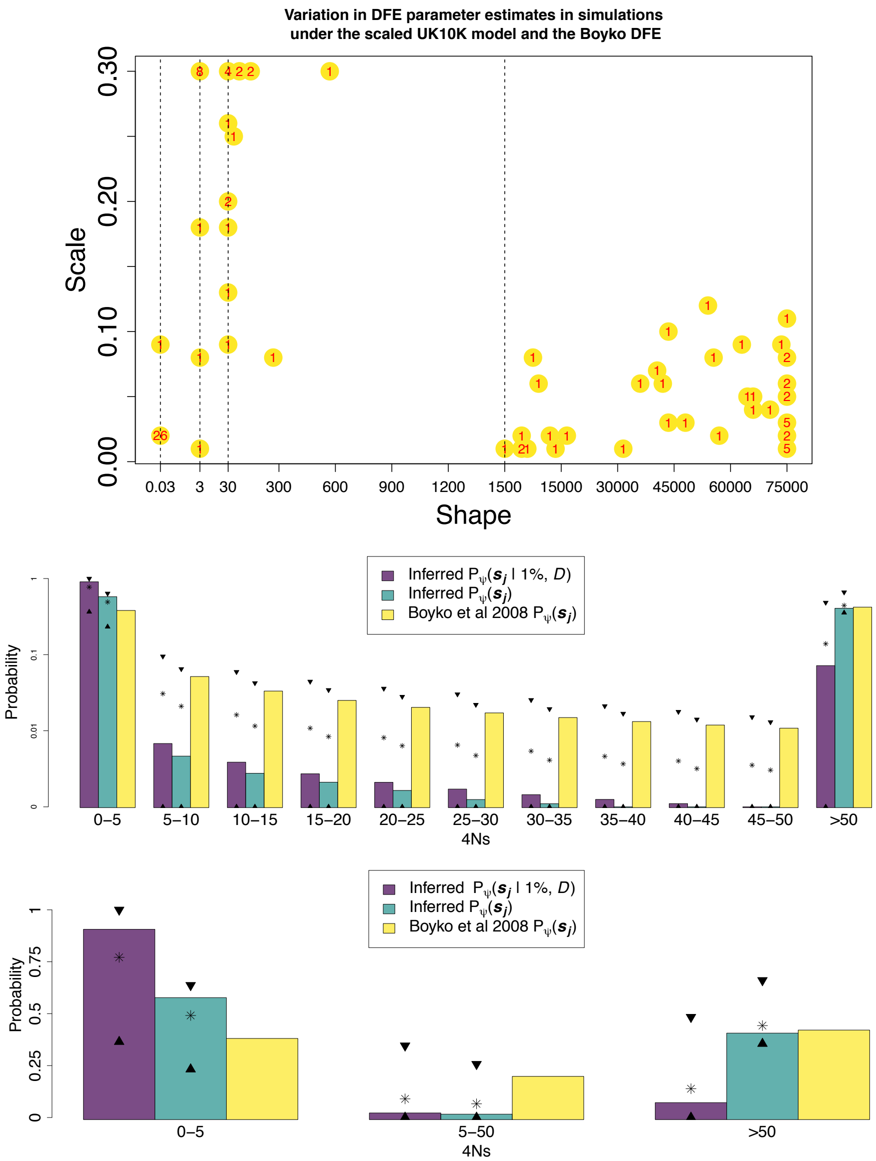


**Supplementary Figure S10.- Variation in estimates of the distribution of fitness effects in 100 sets of simulations of $273$ $1\%$ frequency variants with $72$ haplotypes containing the derived allele in each variant. We simulated these variants under the scaled *UK10K* demographic model and the Boyko et al. 2008 distribution of fitness effects.**
On the top we show the estimated scale and shape parameters of the compound estimated $DFE$ in 100 sets of simulations replicates. The red numbers indicate the number of bootstrap replicates found in each combination of scale and shape parameters.
On the middle panel we compare the estimated values of 'Inferred $P_{\psi}(\boldsymbol{s_j}|f,D)$', 'Inferred $P_{\psi}(\boldsymbol{s_j})$' against the value of 'Boyko et al 2008 $P(\boldsymbol{s_j}|DFE)$'. To calculate 'Inferred $P_{\psi}(\boldsymbol{s_j})$' from the 'Inferred $P_{\psi}(\boldsymbol{s_j}|f,D)$' we used equation (4), calculating $P_{\psi}(f|D)$ from a set of approximately $34.78$ X $10^9$ variants simulated with *PReFerSim* under the scaled *UK10K* demographic model and the Boyko distribution of fitness effects; and $P_{\psi}(f|\boldsymbol{s_j},D)$ from a set of approximately $34.78$ X $10^9$ variants simulated with *PReFerSim* under the scaled *UK10K* demographic model and the

Mouse distribution of fitness effects. The median values of ‘Inferred $P_{\psi}(\boldsymbol{s_j}|f, D)$’ and ‘Inferred $P_{\psi}(\boldsymbol{s_j})$’ across 100 simulations are shown in the bars across each $\boldsymbol{s_j}$ interval. The bar from ‘Boyko et al 2008 $P_{\psi}(\boldsymbol{s_j})$’ is the value inferred for each $\boldsymbol{s_j}$ interval based on the $DFE$ inferred in the Boyko et al 2008 paper. The two triangles shown in each $\boldsymbol{s_j}$ interval denote the $5\%$ and $95\%$ percentile of the ‘Inferred $P_{\psi}(\boldsymbol{s_j})$’ or ‘$P_{\psi}(\boldsymbol{s_j}|f, D)$’ probabilities estimated across the $100$ simulation replicates. The asterisk signs from ‘Inferred $P_{\psi}(\boldsymbol{s_j})$’ and ‘$P_{\psi}(\boldsymbol{s_j}|f, D)$’ probabilities are the mean values calculated from 100 simulation replicates. The actual estimate of ‘Boyko et al 2008 $P_{\psi}(\boldsymbol{s_j})$’ is inside the $5\%$ and $95\%$ percentile of the ‘Inferred $P_{\psi}(\boldsymbol{s_j})$’ values across all $\boldsymbol{s_j}$ intervals. Note that the mean value of ‘Inferred $P_{\psi}(\boldsymbol{s_j})$’ is closer to the true value of ‘Boyko et al 2008 $P_{\psi}(\boldsymbol{s_j})$’ than the median value of ‘Inferred $P_{\psi}(\boldsymbol{s_j})$’ across all $\boldsymbol{s_j}$ intervals .

In the bottom panel, we also compare the estimated values of ‘Inferred $P_{\psi}(\boldsymbol{s_j}|f, D)$’, ‘Inferred $P_{\psi}(\boldsymbol{s_j})$’ against the value of ‘Boyko et al 2008 $P_{\psi}(\boldsymbol{s_j})$’. To calculate ‘Inferred $P_{\psi}(\boldsymbol{s_j}|f, D)$’, ‘Inferred $P_{\psi}(\boldsymbol{s_j})$’ and ‘Boyko et al 2008 $P_{\psi}(\boldsymbol{s_j})$’ for the middle bin $\boldsymbol{s_j} \in [5{,}50)$ we summed the values of those three respective probabilities across eight $\boldsymbol{s_j}$intervals: $\boldsymbol{s_j} \in [5, 10)$, $\boldsymbol{s_j} \in [10, 15)$, $\boldsymbol{s_j} \in [15{,}20)$, $\boldsymbol{s_j} \in [20{,}25)$, $\boldsymbol{s_j} \in [25{,}30)$, $\boldsymbol{s_j} \in [30{,}35)$, $\boldsymbol{s_j} \in [35{,}40)$, $\boldsymbol{s_j} \in [40{,}45)$ and $\boldsymbol{s_j} \in [45{,}50)$. Note that here the y-axis is plotted on a linear scale while in the middle panel we used a logarithmic scale. The two triangles shown in each $\boldsymbol{s_j}$ interval denote the $5\%$ and $95\%$ percentile of the ‘Inferred $P_{\psi}(\boldsymbol{s_j})$’ or ‘$P_{\psi}(\boldsymbol{s_j}|f, D)$’ probabilities estimated across $100$ simulation replicates. The asterisk signs from ‘Inferred $P_{\psi}(\boldsymbol{s_j})$’ and ‘$P_{\psi}(\boldsymbol{s_j}|f, D)$’ probabilities are the mean values calculated from 100 simulation replicates.

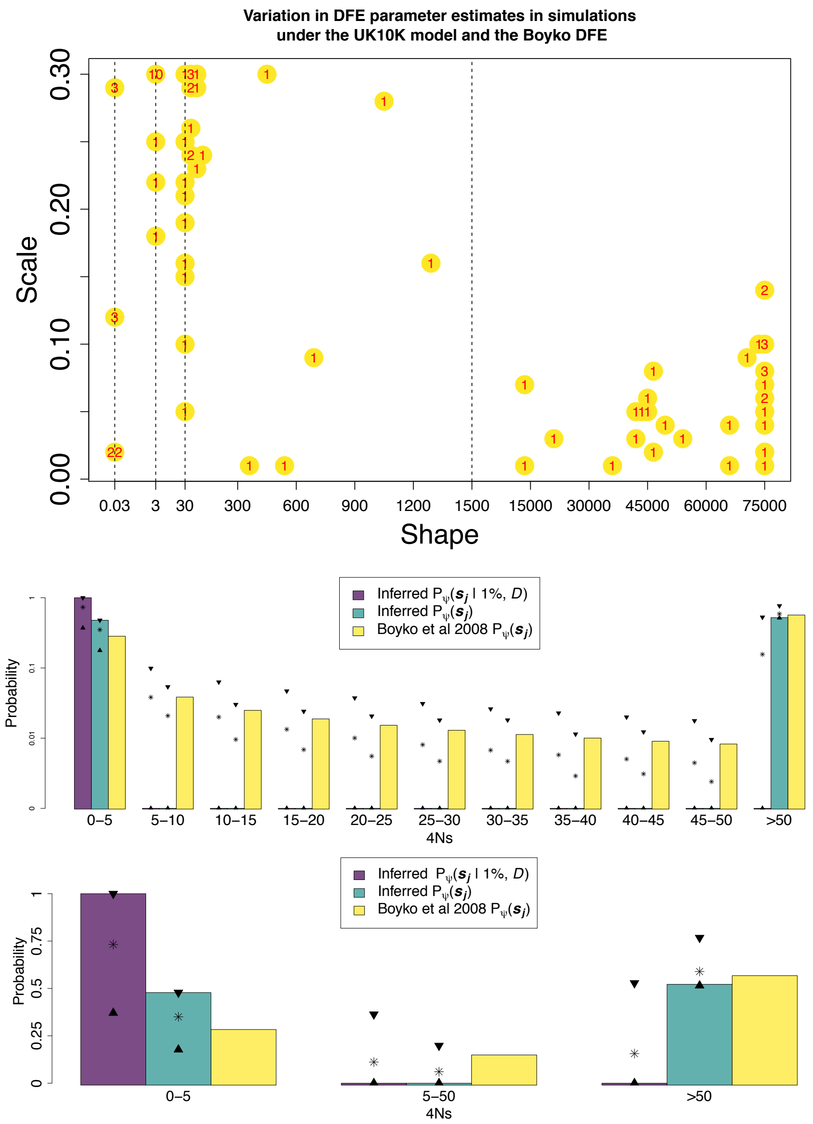


**Supplementary Figure S11.- Variation in estimates of the distribution of fitness effects in 100 sets of simulations of 273 1% frequency variants with 72 haplotypes containing the derived allele in each variant. We simulated these variants under the *UK10K* demographic model and the Boyko et al. 2008 distribution of fitness effects. Here we used the *UK10K* model instead of the scaled *UK10K* model employed in Supplementary Figure S10. The inference of $P_\psi(s_j|f, D)$ was performed using Equation 3 and the scaled *UK10K* demographic model. This mimics the analysis performed in the *UK10K* dataset, where the inferences of *DFE$_f$* were performed using the scaled *UK10K* demographic model.**

On the top we show the estimated scale and shape parameters of the compound estimated $DFE$ in 100 sets of simulations replicates. The red numbers indicate the number of bootstrap replicates found in each combination of scale and shape parameters.

On the middle panel we compare the estimated values of 'Inferred $P_\psi(s_j|f, D)$', 'Inferred $P_\psi(s_j)$' against the value of 'Boyko et al 2008 $P(s_j|DFE)$'. To calculate 'Inferred $P_\psi(s_j)$' from the

‘Inferred $P_\psi(\boldsymbol{s_j}|f,D)$’ we used equation (4), calculating $P_\psi(f|D)$ from a set of approximately $76.16 \text{ X } 10^9$ variants simulated with *PReFerSim* under the *UK10K* demographic model and the Boyko distribution of fitness effects; and $P_\psi(f|\boldsymbol{s_j},D)$ from a set of approximately $76.16 \text{ X } 10^9$ variants simulated with *PReFerSim* under the scaled *UK10K* demographic model and the Mouse distribution of fitness effects. The median values of ‘Inferred $P_\psi(\boldsymbol{s_j}|f,D)$’ and ‘Inferred $P_\psi(\boldsymbol{s_j})$’ across 100 simulations are shown in the bars across each $\boldsymbol{s_j}$ interval. The bar from ‘Boyko et al 2008 $P_\psi(\boldsymbol{s_j})$’ is the value inferred for each $\boldsymbol{s_j}$ interval based on the $DFE$ inferred in the Boyko et al 2008 paper. The two triangles shown in each $\boldsymbol{s_j}$ interval denote the $5\%$ and $95\%$ percentile of the ‘Inferred $P_\psi(\boldsymbol{s_j})$’ or ‘$P_\psi(\boldsymbol{s_j}|f,D)$’ probabilities estimated across the $100$ simulation replicates. The asterisk signs from ‘Inferred $P_\psi(\boldsymbol{s_j})$’ and ‘$P_\psi(\boldsymbol{s_j}|f,D)$’ probabilities are the mean values calculated from 100 simulation replicates. The actual estimate of ‘Boyko et al 2008 $P_\psi(\boldsymbol{s_j})$’ is inside the $5\%$ and $95\%$ percentile of the ‘Inferred $P_\psi(\boldsymbol{s_j})$’ values across all $\boldsymbol{s_j}$ intervals. Note that the mean value of ‘Inferred $P_\psi(\boldsymbol{s_j})$’ is closer to the true value of ‘Boyko et al 2008 $P_\psi(\boldsymbol{s_j})$’ than the median value of ‘Inferred $P_\psi(\boldsymbol{s_j})$’ across all $\boldsymbol{s_j}$ intervals .

In the bottom panel, we also compare the estimated values of ‘Inferred $P_\psi(\boldsymbol{s_j}|f,D)$’, ‘Inferred $P_\psi(\boldsymbol{s_j})$’ against the value of ‘Boyko et al 2008 $P_\psi(\boldsymbol{s_j})$’. To calculate ‘Inferred $P_\psi(\boldsymbol{s_j}|f,D)$’, ‘Inferred $P_\psi(\boldsymbol{s_j})$’ and ‘Boyko et al 2008 $P_\psi(\boldsymbol{s_j})$’ for the middle bin $\boldsymbol{s_j} \in [5,50)$ we summed the values of those three respective probabilities across eight $\boldsymbol{s_j}$intervals: $\boldsymbol{s_j} \in [5,10)$, $\boldsymbol{s_j} \in [10,15)$, $\boldsymbol{s_j} \in [15,20)$, $\boldsymbol{s_j} \in [20,25)$, $\boldsymbol{s_j} \in [25,30)$, $\boldsymbol{s_j} \in [30,35)$, $\boldsymbol{s_j} \in [35,40)$, $\boldsymbol{s_j} \in [40,45)$ and $\boldsymbol{s_j} \in [45,50)$. Note that here the y-axis is plotted on a linear scale while in the middle panel we used a logarithmic scale. The two triangles shown in each $\boldsymbol{s_j}$ interval denote the $5\%$ and $95\%$ percentile of the ‘Inferred $P_\psi(\boldsymbol{s_j})$’ or ‘$P_\psi(\boldsymbol{s_j}|f,D)$’ probabilities estimated across $100$ simulation replicates. The asterisk signs from ‘Inferred $P_\psi(\boldsymbol{s_j})$’ and ‘$P_\psi(\boldsymbol{s_j}|f,D)$’ probabilities are the mean values calculated from 100 simulation replicates.

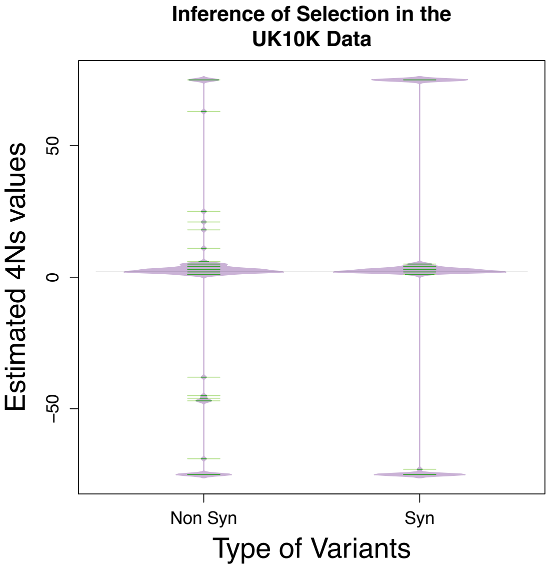


**Supplementary Figure S12.- Variation around the estimated $4Ns$ values using bootstrap replicates.** We performed $100$ bootstrap replicates on the $1\%$ frequency nonsynonymous and synonymous variants. To create the bootstrap replicates, we sampled variants with replacement from our set of synonymous and nonsynonymous variants until we obtained $152$ and $273$ variants, respectively. Then, we estimated the $4Ns$ value for each bootstrap replicate. The point estimates of $4Ns$ in the $\mathbf{273\ 1\%}$ frequency nonsynonymous variants of the *UK10K* dataset and is equal to $4Ns\ = 2$ and $4Ns\ = 3$ for the $152$ $1\%$ frequency synonymous variants. Those point estimates of $4Ns$ are not very different from 0 based on the variation seen in the 100 bootstrap replicates.

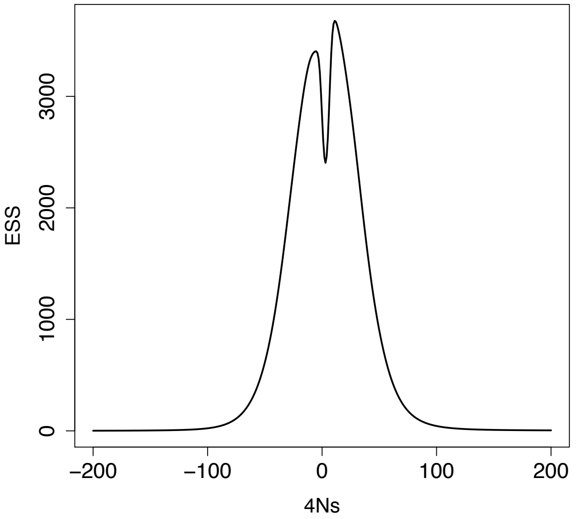


**Supplementary Figure S13.- Effective Sample Sizes (*ESS*) in the inferred scaled *UK10K* demographic scenario of a population expansion using $\mathbf{100,000}$ simulated allele frequency trajectories from our proposal distribution.**

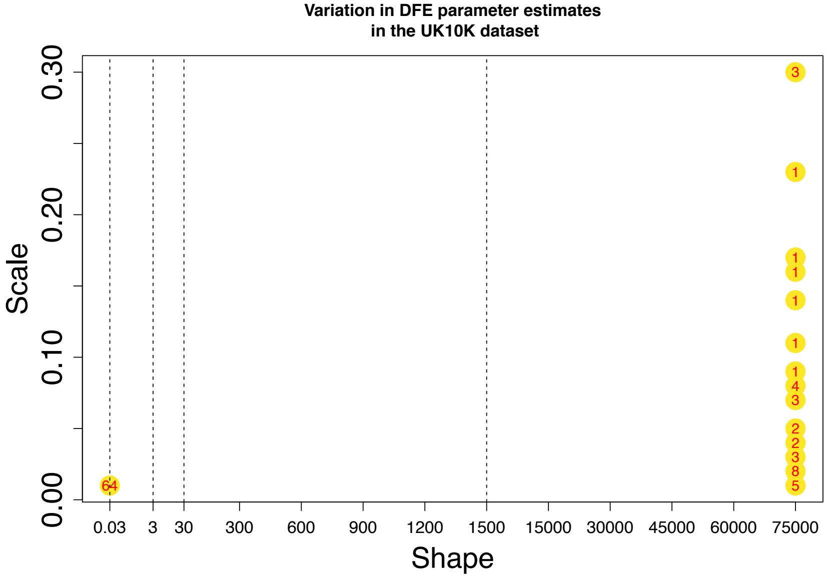


**Supplementary Figure S14.- Variation in estimates of the distribution of fitness effects of 1% frequency nonsynonymous variants in the *UK10K* dataset assessed using a bootstrap approach**.
We created each bootstrap replicate by sampling variants with replacement from our set of nonsynonymous variants until we obtained $273$ variants. A) Estimated scale and shape parameters of the compound distribution in $100$ bootstrap replicates. The red numbers indicate the number of bootstrap replicates found in each combination of scale and shape parameters. We found that estimates of the shape parameter tended to cluster on the edges of the searched parameter values. We did not explore higher values of the shape parameter because we found that our method showed low *ESS*, smaller than $100$, when $4Ns$ was smaller than -$76$ (Supplementary Figure S9).

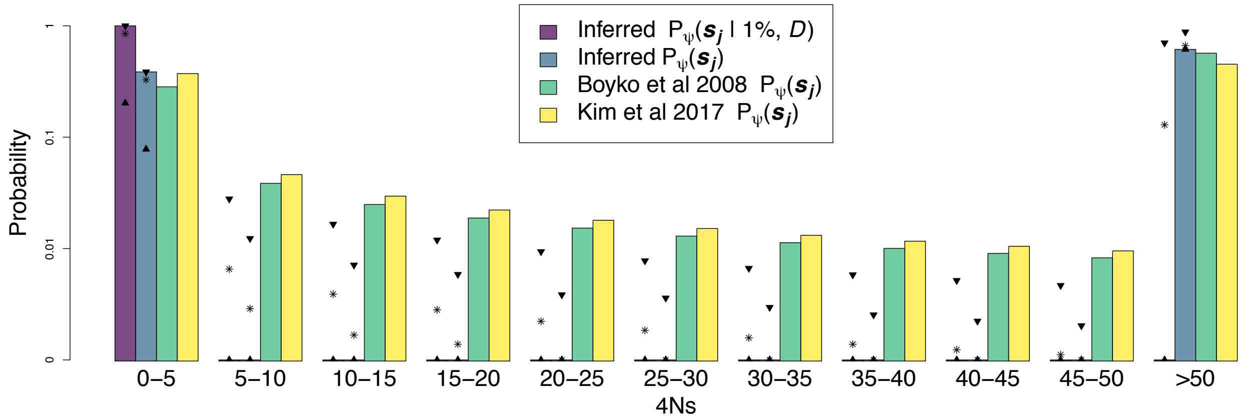


**Supplementary Figure S15.- Inferred distribution of fitness effects of new mutations and 1% frequency deleterious variants in the *UK10K* dataset.** 'Inferred $P_\psi(\boldsymbol{s_j})$' refers to the probability of having a $4Ns$ value in a particular interval $\boldsymbol{s_j}$ given the distribution of fitness effects of new mutations $DFE$. 'Inferred $P_\psi(\boldsymbol{s_j}$ | $1\%,\ D)$' is the probability of having a $4Ns$ value in a particular interval $\boldsymbol{s_j}$ given that the variant has a $1\%$ frequency, and the demographic scenario $D$. The selection coefficient $s$ refers exclusively to the action of deleterious variants in this plot. We compared our inferences with those of Boyko et al. (2008) and Kim et al. (2017). The two triangles shown in each $\boldsymbol{s_j}$ interval denote the upper and lower limit of the $90\%$ bootstrap percentile interval across $100$ bootstrap replicates. The asterisk signs are the mean values for the inferred probabilities $P_\psi(\boldsymbol{s_j}$ | $1\%,\ D)$ and $P_\psi(\boldsymbol{s_j})$ calculated from 100 bootstrap replicates.

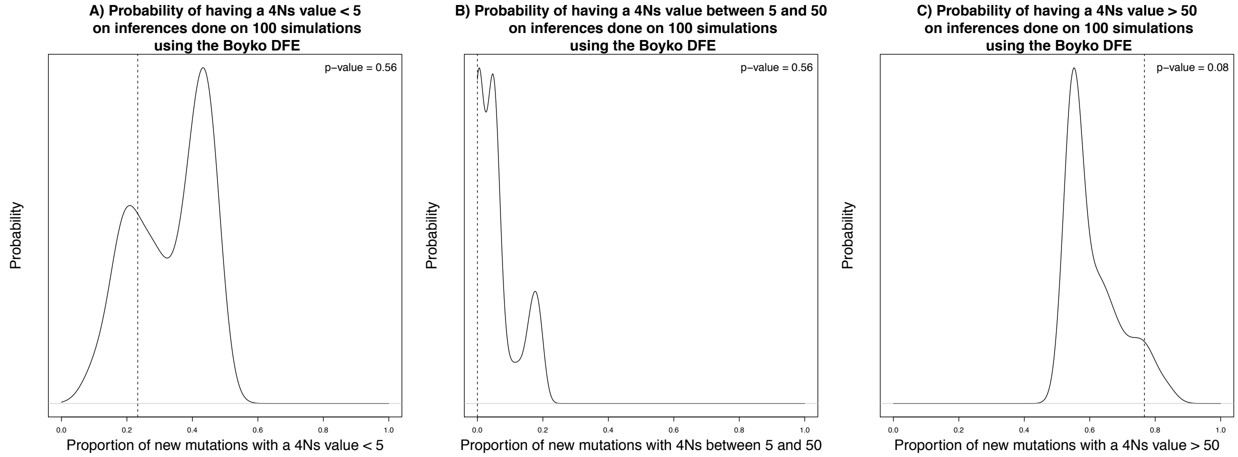


**Supplementary Figure S16.- Probability distributions of the inferred proportion of variants having a A) $4Ns$ value < 5, B) $4Ns$ value between 5 and 50, and c) $4Ns$ value > 50 on 100 simulations done using the Boyko et al. (2008) distribution of fitness effects.** We took the results of the inferred $P(\boldsymbol{s_j}|DFE)$ shown in the bottom plot from Supplementary Figure S11 and created probability density plots for the 3 $\boldsymbol{s_j}$ intervals shown there. We used those results to create null distributions of inferred $P(\boldsymbol{s_j}|DFE)$ values under the assumption that we used the Boyko et al. (2008) $DFE$.
The vertical dashed lines shown in every plot represent the values of $P(\boldsymbol{s_j}|DFE)$ in each $\boldsymbol{s_j}$ bin as calculated from the *UK10K* dataset. We calculated a p-value based on the results from the *UK10K* dataset and the null distributions of $P(\boldsymbol{s_j}|DFE)$ values. Based on the p-values, we cannot reject the null hypothesis that the proportion of variants that have a *4Ns* value in a particular $\boldsymbol{s_j}$ interval in the *UK10K* dataset is different from what is expected under the Boyko distribution of fitness effects.

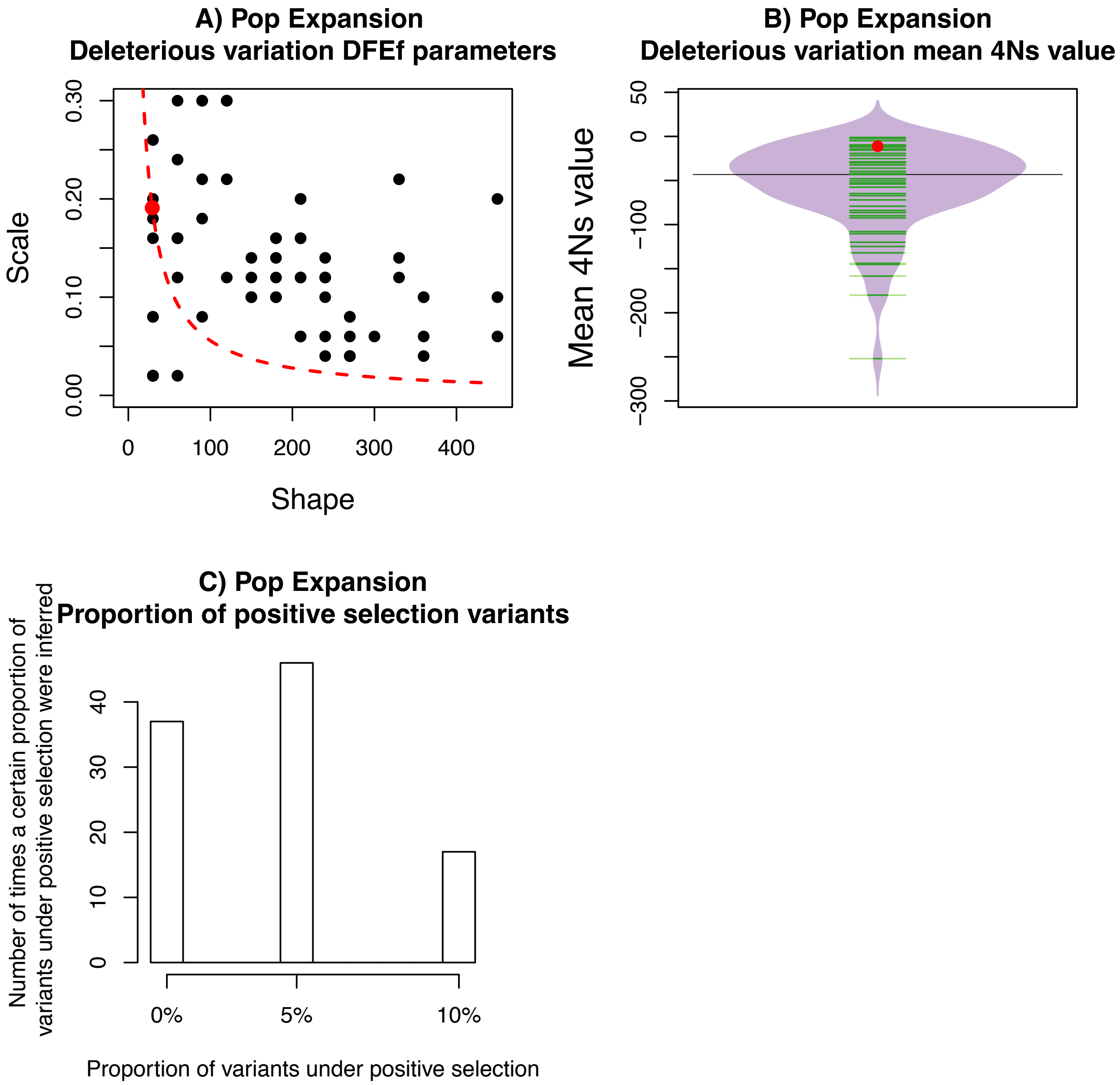


**Supplementary Figure S17.- Estimation of the $DFE_f$ of $1\%$ frequency variants in simulated data when a proportion of variants are under positive selection ($5\%$) and another proportion of the variants are under negative selection under a population expansion model.**
A) The black points denote the estimated $DFE$ parameters. The red dot reflects the actual $DFE$ parameters of a compound distribution that fit the distribution of selection coefficients of $100{,}000$ variants at a $1\%$ frequency. B) Estimated $4Ns$ values. The red dot denotes the actual mean $4Ns$ value in $100{,}000$ variants at a $1\%$ frequency. C) Estimated proportion of variants under positive selection, where the actual value is equal to $5\%$. We assume that the value of $4Ns$ in the variants under positive selection is known, and that we try to estimate the proportion of variants under positive selection along with the parameters of the $DFE_f$ that define the distribution of selection coefficients of $1\%$ variants under negative selection. The grid of explored parameters takes values of $0\%$, $5\%$, $10\%$ for the proportion of variants of positive selection. The grid of

explored parameters for the compound distribution defining the distribution of variants under negative selection goes from $(0.02, 0.04, \ldots, 0.3)$ for the scale parameter and from $(15, 30, \ldots, 450)$ for the shape parameter.

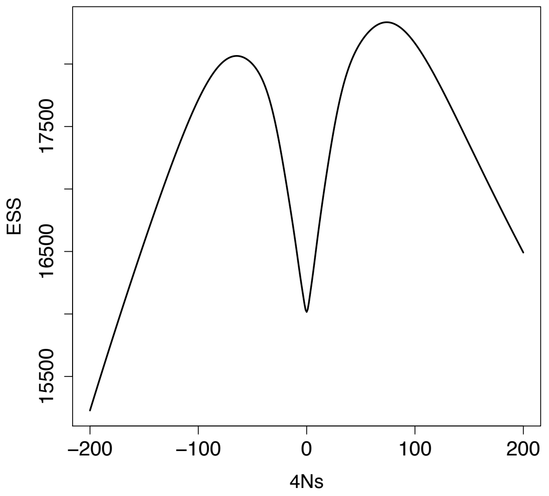


**Supplementary Figure S18.- Effective Sample Sizes (*ESS*) in a demographic scenario of a constant population size using $100,000$ simulated allele frequency trajectories from our proposal distribution.**

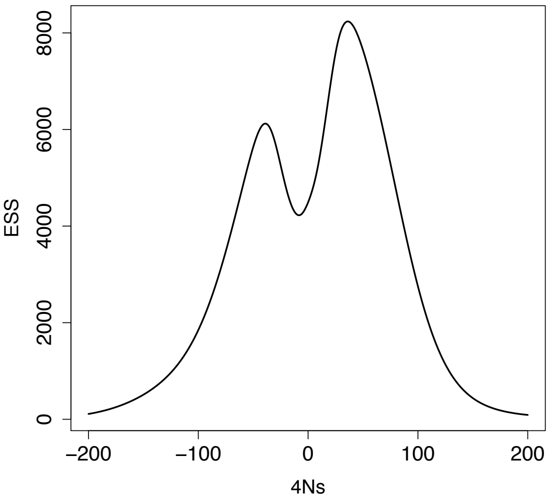


**Supplementary Figure S19.- Effective Sample Sizes (*ESS*) in a demographic scenario of a population expansion using $100,000$ simulated allele frequency trajectories from our proposal distribution.**

**Supplementary Table S1.- Comparison of estimates of the mean allele age of a $1\%$ frequency variant in a demographic scenario of a constant population size ($N = 10,000$).** The theoretical estimates were obtained by Maruyama (1974) and assume that the populations are in the diffusion limit, where $N$ tends to infinity and the value of $4Ns$ tends to a fixed constant. We report an estimate of allele age from forward-in-time simulations using the mean allele age across $10{,}000$ forward-in-time allele frequency trajectories. Our estimate is based on alleles that were sampled at a $1\%$ frequency based on a sample of $4{,}000$ chromosomes. We also report the standard deviation of the allele ages, shown inside parenthesis, for the forward-in-time simulations.

| $4Ns$ | 0 | -50 | 50 | -100 | 100 |
|---|---|---|---|---|---|
| Maruyama's theoretical estimates | 1861 | 654 | 654 | 474 | 474 |
| Forward-in-time simulation estimates based on the sample allele frequency (standard deviation) | 1872.69 (5295.08) | 641.49 (636.20) | 649.00 (635.53) | 462.68 (371.32) | 468.42 (377.05) |

**Supplementary Table S2.- Percentiles of the Inferred '$P_{\psi}(\boldsymbol{s_j})$' or '$P_{\psi}(\boldsymbol{s_j}|f,D)$' probabilities estimated across $100$ simulation replicates for different $s_j$ intervals based on Supplementary Figure S10.**

$P_{\psi}(\boldsymbol{s_j})$

| $\boldsymbol{s_j}$ | 25% and 75% percentile | 10% and 90% percentile | 5% and 95% percentile | 2.5% and 97.5% percentile | Probability from Boyko et al. (2008) scaled by the ancestral population size in the scaled *UK10K* model |
|---|---|---|---|---|---|
| 0-5 | (0.385, 0.637) | (0.286, 0.637) | (0.232, 0.637) | (0.201, 0.637) | 0.381 |
| 5-10 | (0.0, 0.031) | (0.0, 0.060) | (0.0, 0.065) | (0, 0.079) | 0.052 |
| 10-15 | (0.0, 0.015) | (0.0, 0.037) | (0.0, 0.042) | (0.0, 0.051) | 0.033 |
| 15-20 | (0.0, 0.009) | (0.0, 0.029) | (0.0, 0.034) | (0.0, 0.040) | 0.025 |
| 20-25 | (0.0, 0.007) | (0.0, 0.022) | (0.0, 0.028) | (0.0, 0.031) | 0.020 |
| 25-30 | (0.0, 0.006) | (0.0, 0.016) | (0.0, 0.022) | (0.0, 0.023) | 0.017 |
| 30-35 | (0.0, 0.005) | (0.0, 0.014) | (0.0, 0.019) | (0.0, 0.019) | 0.015 |
| 35-40 | (0.0, 0.005) | (0.0, 0.013) | (0.0, 0.017) | (0.0, 0.017) | 0.013 |
| 40-45 | (0.0, 0.004) | (0.0, 0.011) | (0.0, 0.014) | (0.0, 0.015) | 0.012 |
| 45-50 | (0.0, 0.004) | (0.0, 0.011) | (0.0, 0.013) | (0.0, 0.015) | 0.011 |
| >50 | (0.363, 0.510) | (0.363, 0.586) | (0.355, 0.661) | (0.355, 0.688) | 0.421 |

$$P_{\psi}(\boldsymbol{s_j}|f, D)$$

| $s_j$ | 25% and 75% percentile | 10% and 90% percentile | 5% and 95% percentile | 2.5% and 97.5% percentile |
|---|---|---|---|---|
| 0-5 | (0.602, 1.0) | (0.449, 1.0) | (0.365, 1.0) | (0.315, 1.0) |
| 5-10 | (0.0, 0.045) | (0.0, 0.088) | (0.0, 0.095) | (0.0, 0.115) |
| 10-15 | (0.0, 0.021) | (0.0, 0.052) | (0.0, 0.059) | (0.0, 0.071) |
| 15-20 | (0.0, 0.012) | (0.0, 0.038) | (0.0, 0.044) | (0.0, 0.051) |
| 20-25 | (0.0, 0.009) | (0.0, 0.029) | (0.0, 0.036) | (0.0, 0.039) |
| 25-30 | (0.0, 0.008) | (0.0, 0.023) | (0.0, 0.030) | (0.0, 0.031) |
| 30-35 | (0.0, 0.007) | (0.0, 0.018) | (0.0, 0.025) | (0.0, 0.025) |
| 35-40 | (0.0, 0.006) | (0.0, 0.016) | (0.0, 0.021) | (0.0, 0.021) |
| 40-45 | (0.0, 0.005) | (0.0, 0.014) | (0.0, 0.018) | (0.0, 0.019) |
| 45-50 | (0.0, 0.005) | (0.0, 0.013) | (0.0, 0.015) | (0.0, 0.017) |
| >50 | (0.0, 0.255) | (0.0, 0.375) | (0.0, 0.484) | (0.0, 0.534) |

**Supplementary Table S3.- Percentiles of the Inferred '$P_\psi(\boldsymbol{s_j})$' or '$P_\psi(\boldsymbol{s_j}|f, D)$' probabilities estimated across $100$ simulation replicates for different $s_j$ intervals based on Supplementary Figure S11.**

$P_\psi(\boldsymbol{s_j})$

| $\boldsymbol{s_j}$ | 25% and 75% percentile | 10% and 90% percentile | 5% and 95% percentile | 2.5% and 97.5% percentile | Probability from Boyko et al. (2008) scaled by the ancestral population size in the *UK10K* model |
|---|---|---|---|---|---|
| 0-5 | (0.224, 0.452) | (0.175, 0.637) | (0.167, 0.452) | (0.112, 0.452) | 0.283 |
| 5-10 | (0.0, 0.044) | (0.0, 0.048) | (0.0, 0.051) | (0.0, 0.051) | 0.038 |
| 10-15 | (0.0, 0.011) | (0.0, 0.026) | (0.0, 0.029) | (0.0, 0.029) | 0.025 |
| 15-20 | (0.0, 0.006) | (0.0, 0.021) | (0.0, 0.023) | (0.0, 0.023) | 0.019 |
| 20-25 | (0.0, 0.005) | (0.0, 0.017) | (0.0, 0.019) | (0.0, 0.020) | 0.015 |
| 25-30 | (0.0, 0.005) | (0.0, 0.015) | (0.0, 0.017) | (0.0, 0.017) | 0.013 |
| 30-35 | (0.0, 0.005) | (0.0, 0.015) | (0.0, 0.017) | (0.0, 0.018) | 0.011 |
| 35-40 | (0.0, 0.003) | (0.0, 0.009) | (0.0, 0.011) | (0.0, 0.011) | 0.010 |
| 40-45 | (0.0, 0.003) | (0.0, 0.010) | (0.0, 0.012) | (0.0, 0.012) | 0.009 |
| 45-50 | (0.0, 0.003) | (0.0, 0.008) | (0.0, 0.009) | (0.0, 0.009) | 0.008 |
| >50 | (0.548, 0.657) | (0.543, 0.746) | (0.542, 0.781) | (0.542, 0.798) | 0.568 |

$$P_{\psi}(\boldsymbol{s_j}|f, D)$$

| $s_j$ | 25% and 75% percentile | 10% and 90% percentile | 5% and 95% percentile | 2.5% and 97.5% percentile |
|---|---|---|---|---|
| 0-5 | (0.497, 1.0) | (0.388, 1.0) | (0.371, 1.0) | (0.249, 1.0) |
| 5-10 | (0.0, 0.085) | (0.0, 0.093) | (0.0, 0.099) | (0.0, 0.099) |
| 10-15 | (0.0, 0.025) | (0.0, 0.058) | (0.0, 0.063) | (0.0, 0.064) |
| 15-20 | (0.0, 0.013) | (0.0, 0.043) | (0.0, 0.047) | (0.0, 0.048) |
| 20-25 | (0.0, 0.010) | (0.0, 0.033) | (0.0, 0.037) | (0.0, 0.038) |
| 25-30 | (0.0, 0.008) | (0.0, 0.027) | (0.0, 0.031) | (0.0, 0.031) |
| 30-35 | (0.0, 0.007) | (0.0, 0.023) | (0.0, 0.026) | (0.0, 0.027) |
| 35-40 | (0.0, 0.006) | (0.0, 0.020) | (0.0, 0.023) | (0.0, 0.023) |
| 40-45 | (0.0, 0.006) | (0.0, 0.017) | (0.0, 0.020) | (0.0, 0.020) |
| 45-50 | (0.0, 0.005) | (0.0, 0.015) | (0.0, 0.018) | (0.0, 0.018) |
| >50 | (0.0, 0.276) | (0.0, 0.451) | (0.0, 0.528) | (0.0, 0.578) |

**Supplementary Table S4.- Upper and lower limit of different bootstrap percentile intervals of the Inferred '$P_{\psi}(s_j)$' and '$P_{\psi}(s_j|f,D)$' probabilities estimated across 100 bootstrap replicates for different $s_j$ intervals.**

$P_{\psi}(s_j)$

| $s_j$ | 50% bootstrap percentile interval | 80% bootstrap percentile interval | 90% bootstrap percentile interval | 95% bootstrap percentile interval |
|---|---|---|---|---|
| 0-5 | (0.317, 0.386) | (0.175, 0.386) | (0.078, 0.386) | (0.029, 0.386) |
| 5-10 | (0.0, 0.005) | (0.0, 0.011) | (0.0, 0.012) | (0.0, 0.012) |
| 10-15 | (0.0, 0.003) | (0.0, 0.007) | (0.0, 0.007) | (0.0, 0.007) |
| 15-20 | (0.0, 0.002) | (0.0, 0.006) | (0.0, 0.006) | (0.0, 0.006) |
| 20-25 | (0.0, 0.002) | (0.0, 0.004) | (0.0, 0.004) | (0.0, 0.004) |
| 25-30 | (0.0, 0.001) | (0.0, 0.003) | (0.0, 0.004) | (0.0, 0.004) |
| 30-35 | (0.0, 0.001) | (0.0, 0.003) | (0.0, 0.003) | (0.0, 0.003) |
| 35-40 | (0.0, 0.001) | (0.0, 0.002) | (0.0, 0.003) | (0.0, 0.003) |
| 40-45 | (0.0, 0.001) | (0.0, 0.002) | (0.0, 0.002) | (0.0, 0.002) |
| 45-50 | (0.0, 0.001) | (0.0, 0.002) | (0.0, 0.002) | (0.0, 0.002) |
| >50 | (0.614, 0.665) | (0.614, 0.783) | (0.614, 0.880) | (0.614, 0.944) |

$P_{\psi}(\boldsymbol{s_j}|f, D)$

| $\boldsymbol{s_j}$ | 50% bootstrap percentile interval | 80% bootstrap percentile interval | 90% bootstrap percentile interval | 95% bootstrap percentile interval |
|---|---|---|---|---|
| 0-5 | (0.821, 1.0) | (0.453, 1.0) | (0.202, 1.0) | (0.073, 1.0) |
| 5-10 | (0.0, 0.012) | (0.0, 0.026) | (0.0, 0.028) | (0.0, 0.028) |
| 10-15 | (0.0, 0.007) | (0.0, 0.016) | (0.0, 0.017) | (0.0, 0.017) |
| 15-20 | (0.0, 0.005) | (0.0, 0.011) | (0.0, 0.012) | (0.0, 0.012) |
| 20-25 | (0.0, 0.004) | (0.0, 0.009) | (0.0, 0.009) | (0.0, 0.010) |
| 25-30 | (0.0, 0.003) | (0.0, 0.007) | (0.0, 0.008) | (0.0, 0.008) |
| 30-35 | (0.0, 0.003) | (0.0, 0.006) | (0.0, 0.007) | (0.0, 0.007) |
| 35-40 | (0.0, 0.002) | (0.0, 0.005) | (0.0, 0.006) | (0.0, 0.006) |
| 40-45 | (0.0, 0.002) | (0.0, 0.005) | (0.0, 0.005) | (0.0, 0.005) |
| 45-50 | (0.0, 0.002) | (0.0, 0.004) | (0.0, 0.005) | (0.0, 0.005) |
| >50 | (0.0, 0.139) | (0.0, 0.451) | (0.0, 0.703) | (0.0, 0.865) |

**Supplementary Table S5.- Comparison of selection estimates of new mutations using our method and the estimates obtained by Kim et al. (2017) and Boyko et al. (2008).**

The estimates we obtained for the *UK10K* dataset were computed using the median and the 90% bootstrap percentile interval for each of the 4 selection intervals shown below. To do this, we generated 100 bootstrap replicates, where we created each bootstrap replicate by sampling variants with replacement from our set of nonsynonymous variants until we get 273 variants. Then, we estimated the parameters of the compound distribution for each bootstrap replicate. To obtain the proportion of $s$ values in each of the 4 $s$ intervals shown below, we applied equation 4 using our inferred value of $P_{\psi}(\boldsymbol{s_j}|f, D)$ given our estimates of the compound distribution. In the case of $0 \leq s < 2.18 \times 10^{-5}$ we used $\boldsymbol{s_j} \in [0,1)$. We summed the intervals $\boldsymbol{s_j} \in [1,5)$ and $\boldsymbol{s_j} \in [5,10)$ in the case of $2.18 \times 10^{-5} \leq s < 2.18 \times 10^{-4}$. For the bin $2.18 \times 10^{-4} \leq s < 1.09 \times 10^{-3}$, we summed the values of $P_{\psi}(\boldsymbol{s_j})$ across eight bins $\boldsymbol{s_j} \in [10,15)$, $\boldsymbol{s_j} \in [15,20)$, $\boldsymbol{s_j} \in [20,25)$, $\boldsymbol{s_j} \in [25,30)$, $\boldsymbol{s_j} \in [30,35)$, $\boldsymbol{s_j} \in [35,40)$, $\boldsymbol{s_j} \in [40,45)$ and $\boldsymbol{s_j} \in [45,50)$. The probability that $s > 1.09 \times 10^{-3}$ is simply 1 – P($s < 1.09 \times 10^{-3}$).

We compared the estimates of the 90% bootstrap percentile interval with previously obtained estimates. We assume an $N$ value equal to 11,485 for our estimates to obtain the $s$ values in each $\boldsymbol{s_j}$ interval. That $N$ value is equal to the most ancestral population size in the *UK10K* model (see Supplementary Figure S5). We report the median across 100 bootstrap replicates for the *UK10K* model and the numbers in parenthesis correspond to the 90% bootstrap percentile interval.

| | $0 \leq s < 2.18 \times 10^{-5}$ | $2.18 \times 10^{-5} \leq s < 2.18 \times 10^{-4}$ | $2.18 \times 10^{-4} \leq s < 1.09 \times 10^{-3}$ | $s > 1.09 \times 10^{-3}$ |
|---|---|---|---|---|
| Boyko et al. (2008) (African Americans) | 0.211 | 0.111 | 0.111 | 0.568 |
| Kim et al. (2017) (ESP, u = 1.5e-8) | 0.284 | 0.135 | 0.130 | 0.452 |
| *UK10K* | 0.374 (0.063, 0.374) | 1.13e$^{-7}$ (1.13e$^{-7}$, 0.048) | 0.0 (0.0, 0.035) | 0.626 (0.626, 0.859) |
